# *Hammondia hammondi* has a developmental program *in vitro* that mirrors its stringent two host life cycle

**DOI:** 10.1101/148270

**Authors:** Sarah L. Sokol, Abby S. Primack, Sethu C. Nair, Zhee S. Wong, Maiwase Tembo, J.P. Dubey, Shiv K. Verma, Jon P. Boyle

## Abstract

*Hammondia hammondi* is the nearest relative of *Toxoplasma gondii,* but unlike *T. gondii* is obligately heteroxenous. We have compared *H. hammondi* and *T. gondii* development *in vitro* and identified multiple *H. hammondi*-specific growth states. Despite replicating slower than *T. gondii*, *H. hammondi* was resistant to pH-induced tissue cyst formation early after excystation. However, in the absence of stress *H. hammondi* spontaneously converted to a terminally differentiated tissue cyst stage while *T. gondii* did not. Cultured *H. hammondi* could infect new host cells for up to 8 days following excystation, and this period was exploited to generate stably transgenic *H. hammondi*. Coupled with RNAseq analyses, our data clearly show that *H. hammondi* zoites grow as stringently regulated life stages that are fundamentally distinct from *T. gondii* tachyzoites and bradyzoites.

## Introduction

*Toxoplasma gondii*, the causative agent of toxoplasmosis, is a globally ubiquitous apicomplexan parasite capable of infecting all warm-blooded animals, including approximately one-third of the human population (1-4). Although generally asymptomatic in immunocompetent individuals, *T. gondii* is capable of causing disease in the immunocompromised (including HIV/AIDS patients), developing neonates, and occasionally healthy individuals (5-8). The global ubiquity of *T. gondii* is due, at least in part, to its broad host range and ability to persist undetected in an immunocompetent host, recrudescing following immunosuppression (9, 10). These characteristics are lacking in the nearest extant relative of *T. gondii*, *Hammondia hammondi*. Despite sharing >99% of their genes in near perfect synteny (11) and the same definitive host (2), *H. hammondi* is only known to naturally infect rodents, goats, and roe deer (2, 12) where it appears to be relatively avirulent. In addition to its comparatively limited host range, *H. hammondi* has an obligately heteroxenous life cycle (2), where intermediate host stages (i.e., tissue cysts) are only capable of infecting the definitive host, and definitive host stages (i.e., oocysts) can only infect an intermediate host.

In addition to these life cycle differences, these parasite species have *in vitro* behaviors that mirror their differences *in vivo*. Specifically, when *T. gondii* sporozoites (VEG strain; (13)) are used to initiate infections in human host cells, they undergo a period of rapid replication, after which their growth rate slows, and this period is marked by a subpopulation of parasites spontaneously converting to bradyzoite (e.g., cyst-like) stages (13). However *T. gondii* parasites cultivated in this manner still undergo multiple cycles of replication, egress and reinvasion, and can be propagated indefinitely (13). In contrast, when *H. hammondi* sporozoites are put into tissue culture they replicate more slowly than *T. gondii* (replication predicted to occur once every 24 hours as opposed to once every 6 hours for *T. gondii*) (13, 14), have not been observed to be capable of subculture (either via passage of culture supernatants or after scraping and trypsinization) (15) (14)), and ultimately form bradyzoite cyst-like stages that are infectious to the definitive (but not the rodent intermediate) host (15) (14).

While the molecular determinants that control these dramatic phenotypic differences are unknown, the fact life cycle restriction in *H. hammondi* occurs *in vitro* provides a unique opportunity to identify key aspects of *H. hammondi* biology that underlie its restrictive life cycle and, by contrast, allow for elucidation of the cellular and molecular differences in *T. gondii* that underlie its comparatively flexible life cycle. It is important to note that with respect to other apicomplexans (including *N. caninum* and *Plasmodium* spp.), it is the *T. gondii* life cycle that is atypical; the majority of apicomplexans are obligately heteroxenous. One hypothesis is that *T. gondii* has been released from an ancestral state characterized by a restrictive life cycle and that this ancestral phenotype was more similar to that of *H. hammondi*.

Here, we have characterized *in vitro* excystation, invasion, and replication rate of *T. gondii* (VEG strain; (16, 17)) and *H. hammondi* (HhCatEth1 and HhCatAmer (12, 18)) from sporozoite-initiated infections, and our work represents the most thorough head-to-head comparison of the intermediate host life stages of these genetically similar but phenotypically distinct parasite species. We have also compared the timing and frequency of *in vitro* differentiation and uncovered novel aspects that further illustrate the rigidity of the *H. hammondi* life cycle. Finally, we report the first ever transcriptome from replicating *H. hammondi*, and identified a transcriptional profile that reflects its slow growth and propensity for cyst formation, which fully differentiates it from *T. gondii*. Using this information, we were able to successfully subculture *H. hammondi in vitro* and in mice, as well as define the precise window of infectivity and replicative capacity. With these limits in mind, we generated the first ever transgenic *H. hammondi,* opening up the door to future molecular and genetic studies in this poorly understood parasite species.

## Results

### *T. gondii and H. hammondi* have similar ability to infect host cells

While *T. gondii* oocysts and sporozoites have been studied extensively for their ability to cause disease in intermediate hosts (13, 19, 20), very little is known about the biology of *H. hammondi* oocysts and sporozoites. We compared sporozoite yield and infectivity between *T. gondii* and *H. hammondi* using oocyst preparations derived from multiple cat infections and consistently found that the yield of sporozoites from HhCatEth1 was lower than that from TgVEG oocysts. Average HhCatEth1 sporozoite yield (Mean=5.0 SD=1.2, n=21) was significantly lower than *TgVEG* yield (Mean=11.6 SD=2.13, n=10) (Fig 1A; P=0.02). While excystation yields were lower for HhCatEth1, a linear regression demonstrates no significant relationship between sporozoite yield as a product of oocyst age for HhCatEth1 (*P* =0.76, R^2^=0.01) or TgVEG (*P* =0.63, R^2^=0.03; Fig 1B). Given that sporozoite yields were different between species, we determined if there were differences in sporozoite infectivity by using vacuole formation as a proxy. Interestingly, TgVEG and HhCatEth1 sporozoites had similar infectivity rates in HFFs, defined as the number of vacuoles counted as a fraction of number of vacuoles expected (HhCatEth1 0.12 +/- 0.02 N=2; TgVEG; 0.10 +/- 0.04 N=2 *P*=0.64).

**Fig 1.**
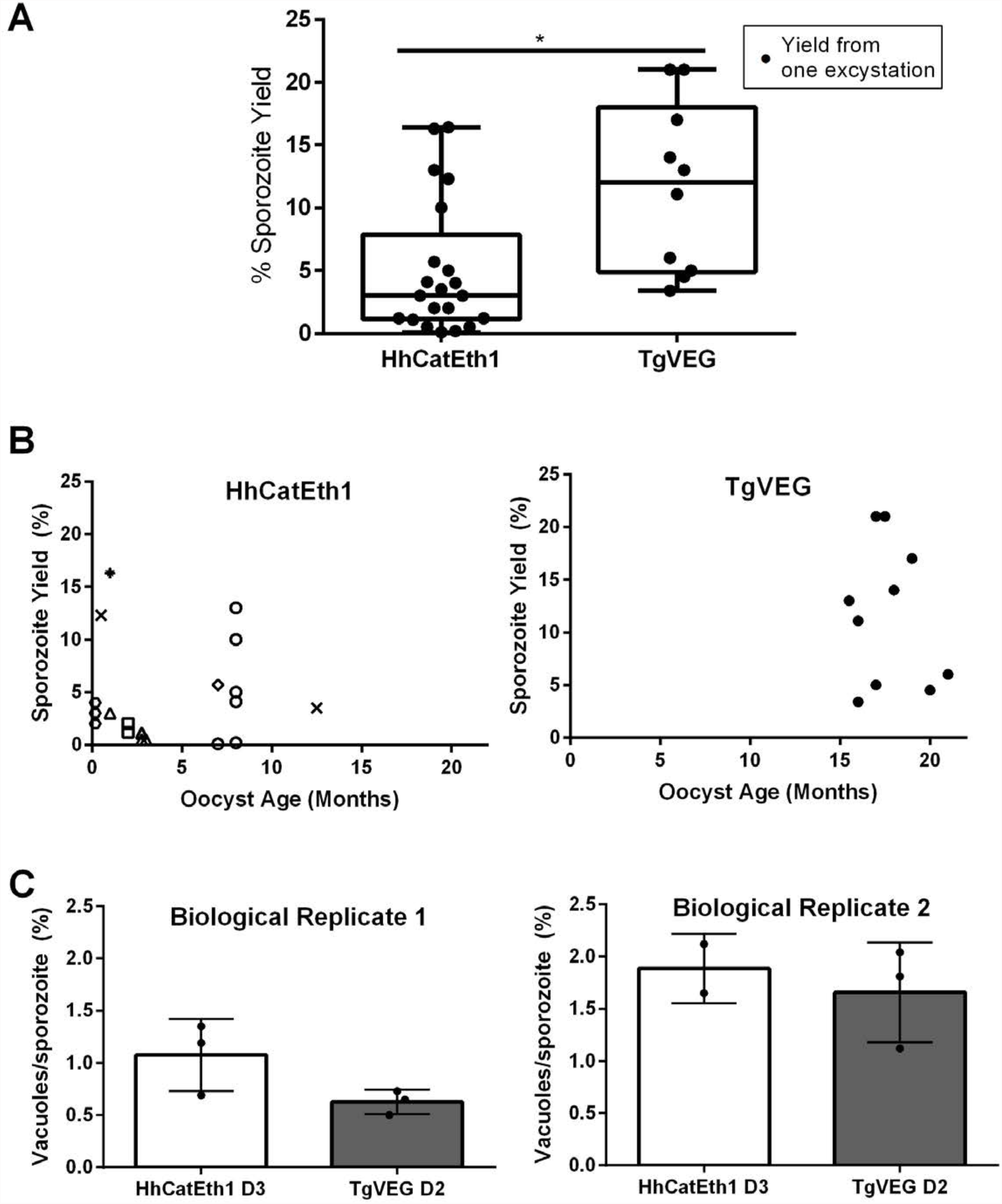
*T. gondii* and *H. hammondi* maintain similar infection rates, but *T. gondii* excysts at significantly higher rates. A) The average sporozoite yield from an excystation of HhCatEth1 is significantly lower (5.0±1.2%) than the average yield of TgVEG sporozoites (11.6±2.1%). Significance determined via unpaired T-test with Welch’s correction, *P*=0.02. B) HhCatEth1 and TgVEG sporozoite yields from oocyst preps of different ages. Sporozoite yield is not significantly influenced by age of oocysts. Significance determined by linear regression: HhCatEth1 *P*=0.76, TgVEG *P*=0.63. Each shape represents one yield from an independent biological prep. C) There is no significant difference in sporozoite viability between HhCatEth1 and TgVEG. Graphs demonstrate the *in vitro* percent viability (measured by the total number of vacuoles/vacuoles expected) of HhCatEth1 and TgVEG sporozoites post excystation. Bars show the mean and SD of two biological replicates (HhCatEth1 0.1202±0.0243, TgVEG 0.1035±0.0346) and there was no significance in viability between species, determined via unpaired T-test with Welch’s correction, *P*=0.6384.

### *H. hammondi* displays differences in *in vitro* replication rate when compared to *T. gondii.*

Although *H. hammondi* is capable of limited *in vitro* growth, the dynamics of these infections with regards to replication rate has yet to be determined. We performed two *in vitro* assays with different cat-derived oocyst preparations to quantify the replication rate of *H. hammondi* (HhCatEth1) and *T. gondii* (TgVEG) sporozoites. As expected from previous work, these assays showed that HhCatEth1 has a slower division rate than TgVEG (Fig 2A). In both replicates, we observed significant growth differences between TgVEG during all 3 days of quantification (** *P*= <0.01, **** *P*= <0.0001; Fig 2B,C); however, we only observed significant growth differences between day 1 and day 3 in one replicate (** *P*= <0.01; Fig 2C). Replication proceeded in TgVEG with an division rate of 2-3x that of HhCatEth1, based on median vacuole size (Table 1, Fig 2D and E). Overall, we’ve calculated the maximum replication rate of TgVEG to be once every 9.6 hours (Day 2) and once every 18 hours for HhCathEth1 (Day 3).

**Fig 2.**
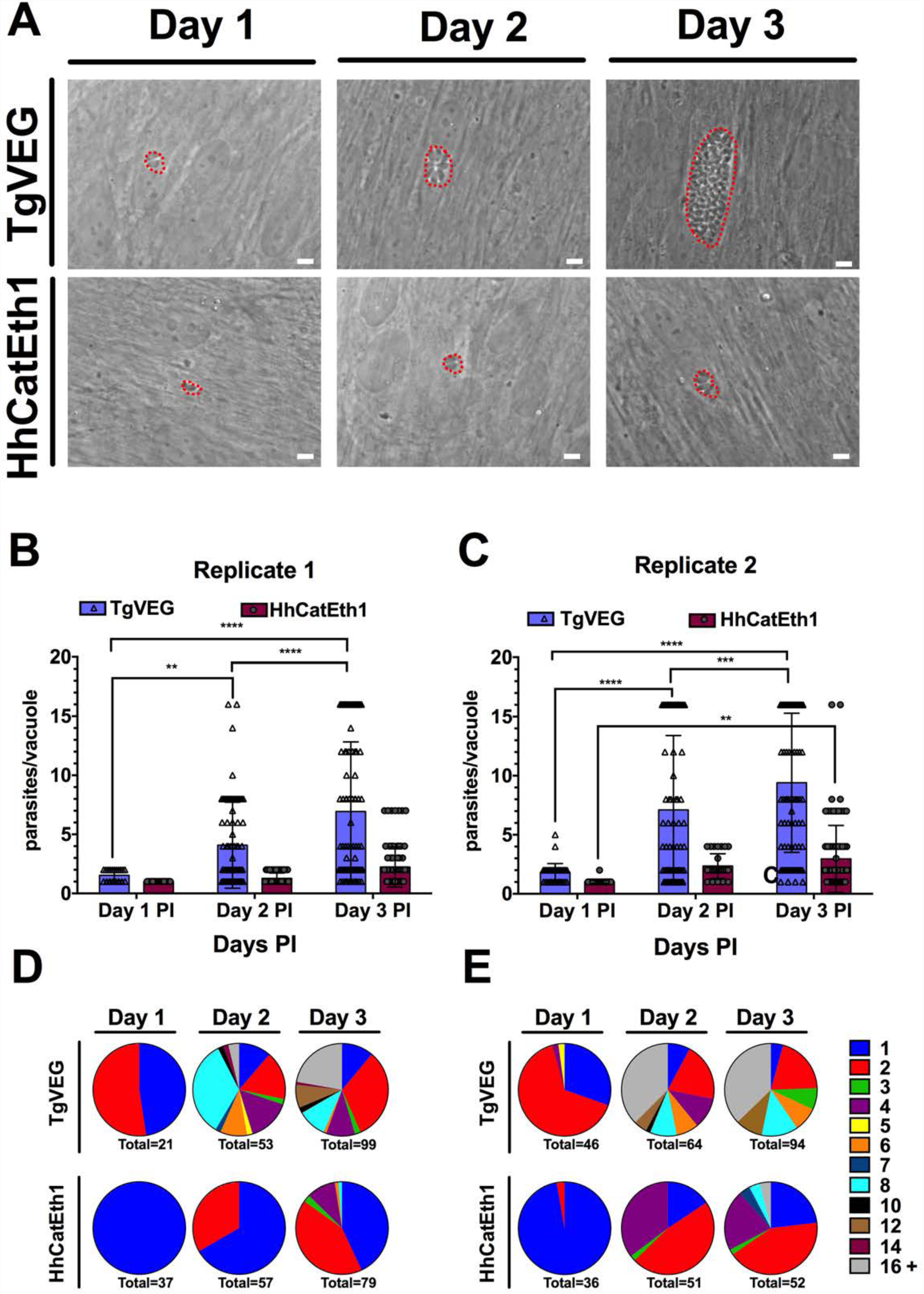
*H. hammondi* replicates at a decreased rate compared to *T. gondii.* A) Confluent monolayers of HFFs were infected with either HhCatEth1 or TgVEG sporozoites at an MOI of 0.5 and visualized over 3 days. While TgVEG sporozoites rapidly divided and produced large vacuoles by 3 DPI, *H. hammondi* did not. B and C) Confluent monolayers of human foreskin fibroblasts (HFFs) were infected with either HhCatEth1 or TgVEG sporozoites, obtained via *in vitro* excystation at an MOI of 0.5. Infected monolayers were fixed at 1, 2, and 3 DPI and assayed for the number of parasites observed in each vacuole for infection with *T. gondii and H. hammondi* for a total of 2 replicates, Replicate 1 (B) and Replicate 2 (C). Vacuoles containing more than 16 parasites were binned at 16 as it was defined as the limit of confident detection. Bars represent mean ± SD. For both Biological Replicate 1 & 2, there was a significant difference between the number of parasites per vacuole on Days 1, 2, &3 PI for TgVEG. For Biological replicate 1, there was no significant difference between the number of parasites per vacuole for HhCatEth1 on Days 1, 2 & 3 PI. For Biological Replicate 2, there was a significant difference in the number of parasites per vacuole between 1 and 3 DPI for HhCatEth1. Statistical significance was determined with a 2-Way ANOVA and Tukey’s multiple comparison test. D and E) Pie chart representing proportion of *T. gondii and H. hammondi* vacuoles containing different number of parasites after 1, 2, and 3 DPI for Replicate 1 (D) and Replicate 2 (E). Vacuoles containing 16 or more vacuoles were donated as 16+.

**Table 1:** Median and maximum parasites per vacuole during 1, 2, and 3 DPI for TgVEG and HhCatEth1.

|  | Day 1 | Day 2 | Day 3 |
| --- | --- | --- | --- |
|  | Median, Maximum | Median, Maximum | Median, Maximum |
| HhCatEth1 Replicate 1 | 1, 1 | 1, 2 | 2, 4 |
| HhCatEth1 Replicate 2 | 1, 2 | 2, 4 | 2, 16 |
| **TgVEG**<br>**Replicate 1** | 2, 2 | 2, 16 | 4, 16+ |
| **TgVEG**<br>**Replicate 2** | 2, 4 | 4, 16+ | 8, 16+ |

### Both *T. gondii* and *H. hammondi* spontaneously form tissue cysts, but do so with different dynamics and efficacy

*T. gondii* and *H. hammondi* are both known to spontaneously differentiate into bradyzoites and form tissue cysts during *in vitro* culture(2). *T. gondii* sporozoite derived infections spontaneous express the bradyzoite antigen 1 (BAG1) after 6 days of growth (13). Yet, the temporal dynamics of differentiation and tissue cyst formation remain unknown in *H. hammondi*. To characterize the efficiency of tissue cyst formation, we infected HFFs with TgVEG and HhCatEth1 sporozoites and stained them with *Dolichos biflorus* Agglutinin (DBA) at 4 and 15 DPI to identify tissue cysts. All vacuoles in TgVEG and HhCatEth1 were DBA-negative at 4 DPI. At 15 DPI, 15% (34/355) of TgVEG vacuoles were DBA-positive, ~80% (152/182) of *H. hammondi* vacuoles were DBA-positive (P<0.0001; Fig 3A, B). To more precisely determine the temporal dynamics of tissue cyst formation, we performed excystations and infected HFFs with TgVEG (Fig 3C) or HhCatAmer (Fig 3D) sporozoites and quantified DBA-positive vacuole formation over the course of 23 days. As expected, we found that both TgVEG and HhCatAmer spontaneously formed DBA-positive tissue cysts (as early as 8 and 12 DPI, respectively) but by 23 DPI 100% of all HhCatAmer vacuoles were DBA-positive, while DBA-positive TgVEG vacuoles never exceeded 20% (Fig 3C,D). These data show that *H. hammondi* undergoes a much more rigid developmental program during *in vitro* cultivation compared to *T. gondii*, resulting in 100% tissue cyst conversion.

**Fig 3.**
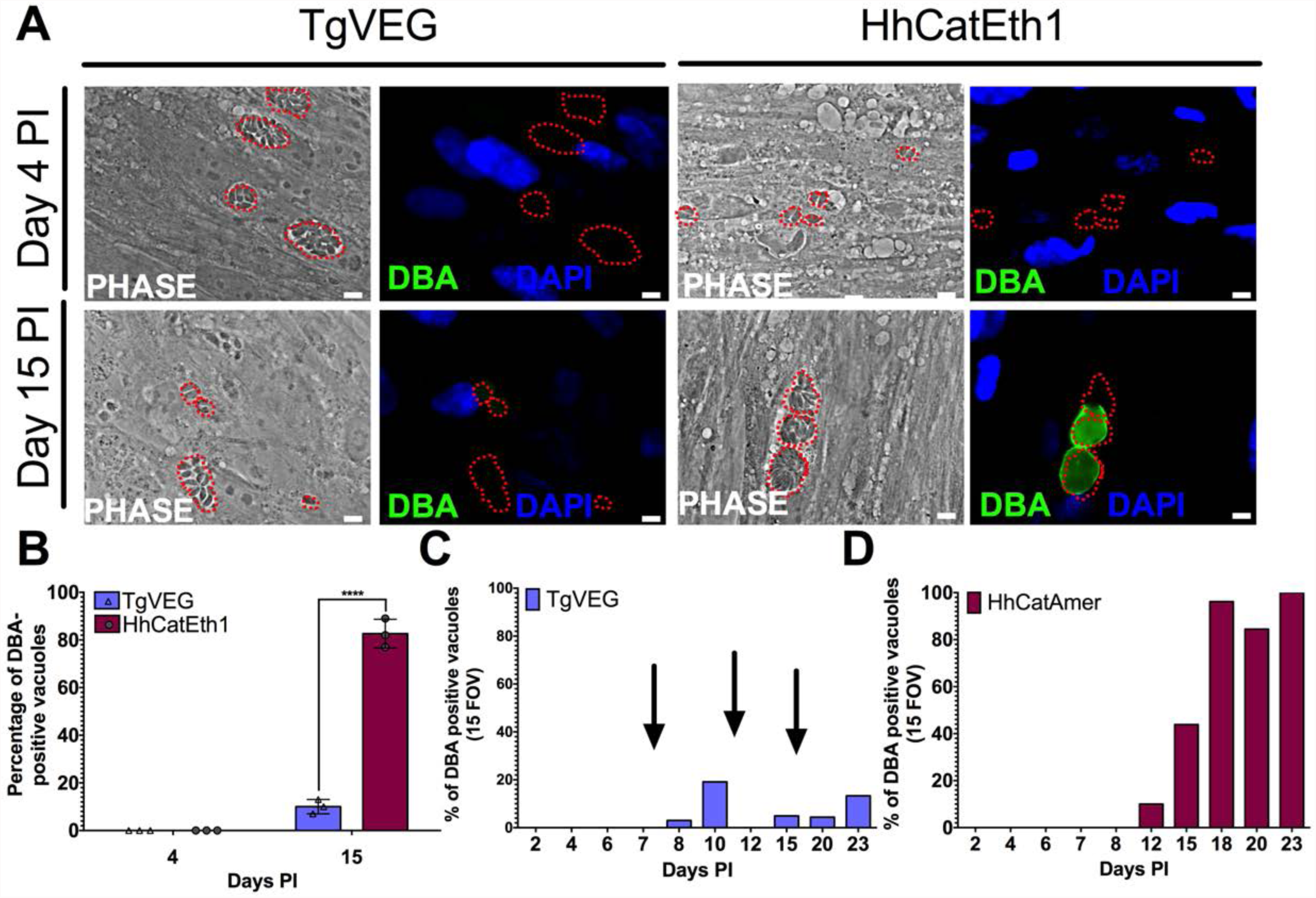
*T. gondii* & *H. hammondi* spontaneously form tissue cysts *in vitro*, but do so with different dynamics. A) Representative images of cells infected with *T. gondii* (TgVEG) and *H. hammondi* (HhCatEth1) sporozoites that were fixed and stained with DBA after 4 and 15 DPI. Scale bar represents 5μm. B) Quantification of *T. gondii* and *H. hammondi* infection described in A demonstrating that after 4 days, neither *H. hammondi* or *T. gondii* spontaneously form tissue cysts, but both parasite species form tissue cysts after 15 DPI. However, the percentage of *H. hammondi* tissue cysts is significantly increased after 15 DPI when compared to *T. gondii.* Numbers above bars indicate number of DBA+ vacuoles out of total vacuoles quantified. Statistical significance was determined using a 2-way ANOVA with Sidak’s multiple comparison test. (****=P<0.0001). C and D) Average percentage of DBA positive vacuoles in 15 FOV over a course of a 23 day infection with *T. gondii* (TgVEG) & *H. hammondi* (HhCatAmer). Arrows in (C) indicate when *T. gondii* was passed onto new host cells to prevent complete lysis. D) *H. hammondi* form tissue cysts after 12 DPI. The percentage of DBA positive vacuoles increases until all vacuoles are DBA positive at 23 DPI. C) *T. gondii* forms tissue cysts at 8 DPI, however, the percentage of DBA positive vacuoles does not reach 100% during the 23 day infection.

### *H. hammondi* is resistant to *in vitro* conditions that induce cyst formation in *T. gondii.*

While *T. gondii* can spontaneously form tissue cysts following an *in vitro* sporozoite infection, it can also be chemically induced to form bradyzoites using treatments such as alkaline pH (13). The responsiveness of *H. hammondi* to these stressors is not known. To determine if we can induce tissue cyst formation in *H. hammondi* prior to its natural progression to a 100% DBA-positive population, we exposed *T. gondii* and *H. hammondi* sporozoites (TgVEG and HhCatAmer) to high pH bradyzoite induction media during 2-4 DPI. Based on DBA staining, we found that while 35% of TgVEG vacuoles grown in bradyzoite induction media for 2 days were DBA-positive (104 out of 297), we could not find a single DBA-positive HhCatAmer vacuole (out of 59 vacuoles; Fig 4A,B). Although a decrease in parasite replication is linked to bradyzoite formation (13) (21), we did not observe a significant difference in the number of parasites per vacuole between TgVEG cultured at pH 7.2 and TgVEG at pH 8.2 (P=0.63) or in HhCatAmer cultured at pH 7.2 and HhCatAmer at pH 8.2 (P=1.00). We did, however, observe an expected significant difference in the number of parasites per vacuole between TgVEG and HhCatAmer in both conditions (** P= 0.0045; **** P= < 0.0001) consistent with the fact that *T. gondii* and *H. hammondi* replicate at different rates (Fig 2). The pH resistance of *H. hammondi* is yet another trait that distinguishes this parasite from *T. gondii*, and provides further evidence that the *H. hammondi* developmental program is hard-wired and very difficult to disrupt.

**Fig 4.**
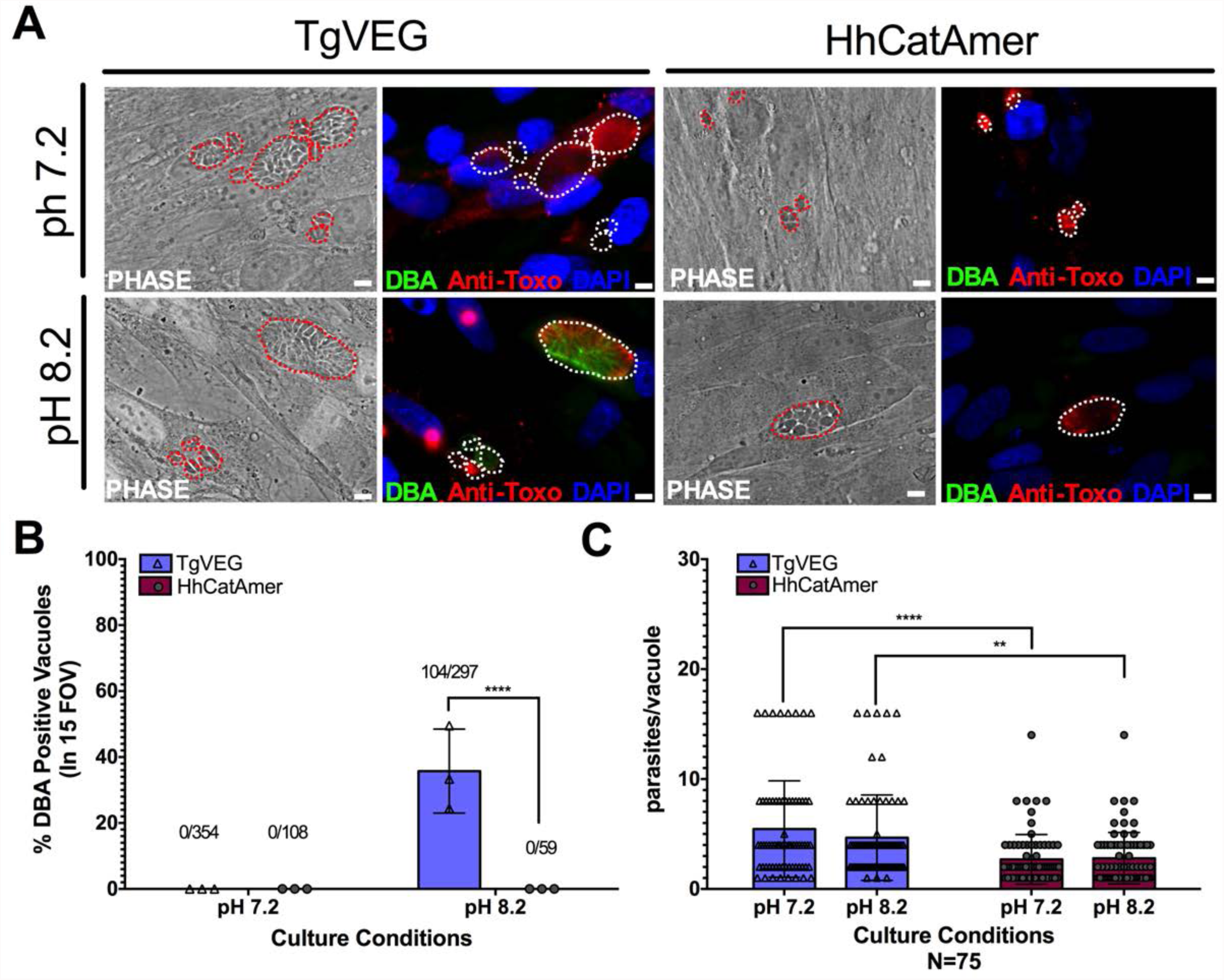
Conditions that induced *T. gondii* tissue cyst formation *in vitro* do not induce tissue cyst formation for *H. hammondi*. A) Representative images of cells infected with *T. gondii* (TgVEG) and *H. hammondi* (HhCatAmer) sporozoites that were grown for 2 days in pH 7.2 media and switched to pH 8.2 media (bradyzoite induction conditions) for 2 days, fixed and stained with an Anti-*Toxoplasma gondii*/*Hammondia hammondi* antibody and DBA. Scale bar represents 5μm. B) Quantification of *T. gondii* and *H. hammondi* tissue cyst formation described in A demonstrating that *H. hammondi* does not form tissue cysts when exposed to conditions that promote tissue cyst formation in *T. gondii.* Number above bars indicated number of vacuoles quantified for each condition. Statistical significance was determined using a 2-way ANOVA with Tukey’s multiple comparison test. (****=P<0.0001). C) Number of parasites per vacuole for both *T. gondii* & *H. hammondi* grown at pH 7.2 and pH 8.2. *T. gondii* has significantly more parasites per vacuole than *H. hammondi* at both pH 7.2 and pH 8.2. There is no statistically significant difference between either *T. gondii* or *H. hammondi* between pH 7.2 and pH 8.2. Statistical significance was determined using a 2-way ANOVA with Sidak’s multiple comparison test. (**=P<0.01 & ****=P<0.0001).

### *H. hammondi* can be subcultured successfully *in vitro*

A key distinguishing feature of *H. hammondi* compared to all *Toxoplasma* strains studied to date is the inability to culture the parasites indefinitely *in vitro*. A consistent observation has been that when *H. hammondi* sporozoites are used to infect host cells (from multiple organisms including humans, cattle, and non-human primates), after a brief time in culture parasites do not retain the ability to infect new host cells (14, 15). To more precisely define the timing of this phenotype, we subcultured *H. hammondi* sporozoites (by needle passage) at multiple times post-excystation (Fig 5A-B). We found that when *H. hammondi* zoites were mechanically released from their host cells, they were capable of infecting and replicating within new host cells for up to 8 days post-excystation (Fig 5C-F), after which we were unable to observe any evidence of parasite replication. In agreement with previous studies (2, 14, 15) we also observed that the number of visible *H. hammondi* vacuoles seems to disappear the longer the parasites were grown in culture. Additionally, we observed that vacuoles began to disappear after 7 DPI (Fig 5C-E). Vacuoles grew in size over this over 13 day incubation period, while new vacuoles derived from lysing and re-invading parasites (which would have comparatively smaller sizes) were never observed after 10 DPI (S1 Fig). These data support previous work showing that *H. hammondi* cannot be subcultured with high efficiency after 7 days in culture (14, 15), but identify a previously unknown window where this parasite can be effectively subcultured. These data also show that parasites emerging from lysed vacuoles at the later stages of incubation are impaired in their ability to re-invade host cells, a phenotype that is again highly distinct from *T. gondii*.

**Fig 5.**
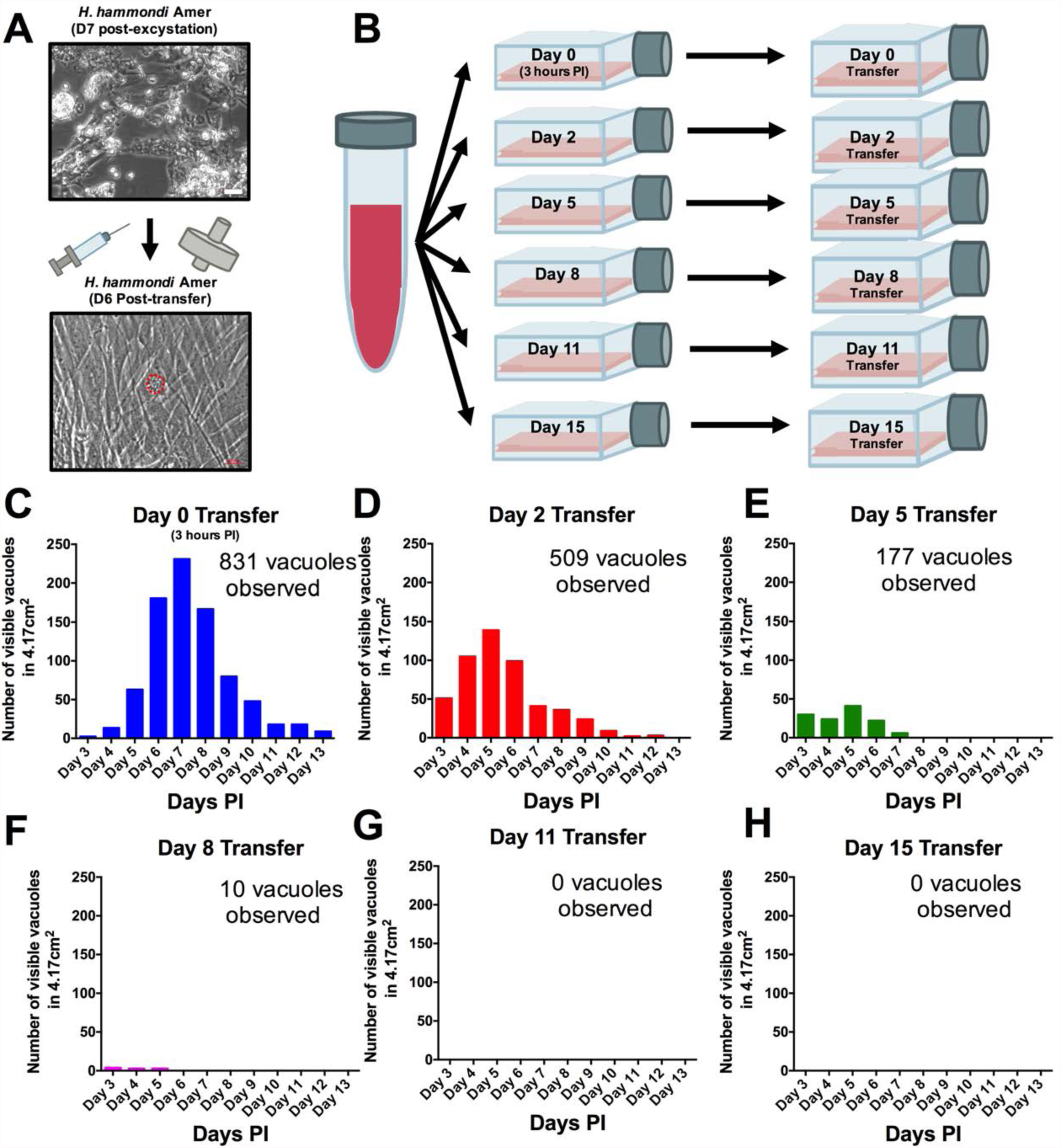
*H. hammondi* can be successfully subcultured *in vitro* for a limited period of time. A) Monolayers containing HhCatAmer with oocyst debris was needle passaged, filtered, and used to infect a confluent monolayer of HFFs seven days post-excystation. Vacuoles were observed in subcultured monolayers after 6 DPI. B-H) Confluent monolayers of HFFs were infected with HhCatAmer sporozoites at an MOI of ~2. (2,812,500 sporozoites). After a three hour incubation at 37° C, the Day 0 transfer flask was scraped, needle passaged, filtered, and transferred to a new host cells. This process was repeated after 2, 5, 8, and 15 days of growth. Each flask was monitored daily for the number (C-H) for 13 DPI, or until visible vacuoles were no longer detected.The number of visible vacuoles increased in number for the first week post-excystation then began to decrease. Replicative capacity was greatest for sporozoites transferred to new host cells at Day 0 (C) and decreased during each subsequent passage. (D-H).

### *H. hammondi* zoites from early *in vitro* culture are infective to mice

Numerous reports (2, 14, 15, 22) have demonstrated that *H. hammondi* grown *in vitro* cannot be used to infect another intermediate host, but our data in Fig 5 show that parasites can be subcultured during first 8 days post-excystation. Here, we aimed to determine if *H. hammondi* cultivated for a short time *in vitro* could also be used to infect mice. HhCatAmer or HhCatEth1 sporulated oocysts were excysted and grown in HFFs for 4 days, at which time monolayers were scraped, syringe lysed, and filtered. We infected mice intraperitoneally with 50,000 zoites of either HhCatAmer (4 mice) or HhCatEth1 (1 mouse) and harvested spleens, peritoneal lavage fluid, and cells (PECs) on 4 (3 mice) or 9 (2 mice) DPI. All 5 spleen samples and 1 PEC sample had detectable *H. hammondi* DNA based on PCR using *H. hammondi* specific primers (Fig 6A), showing that early cultures of *H. hammondi* maintained mouse infectivity.

**Fig 6.**
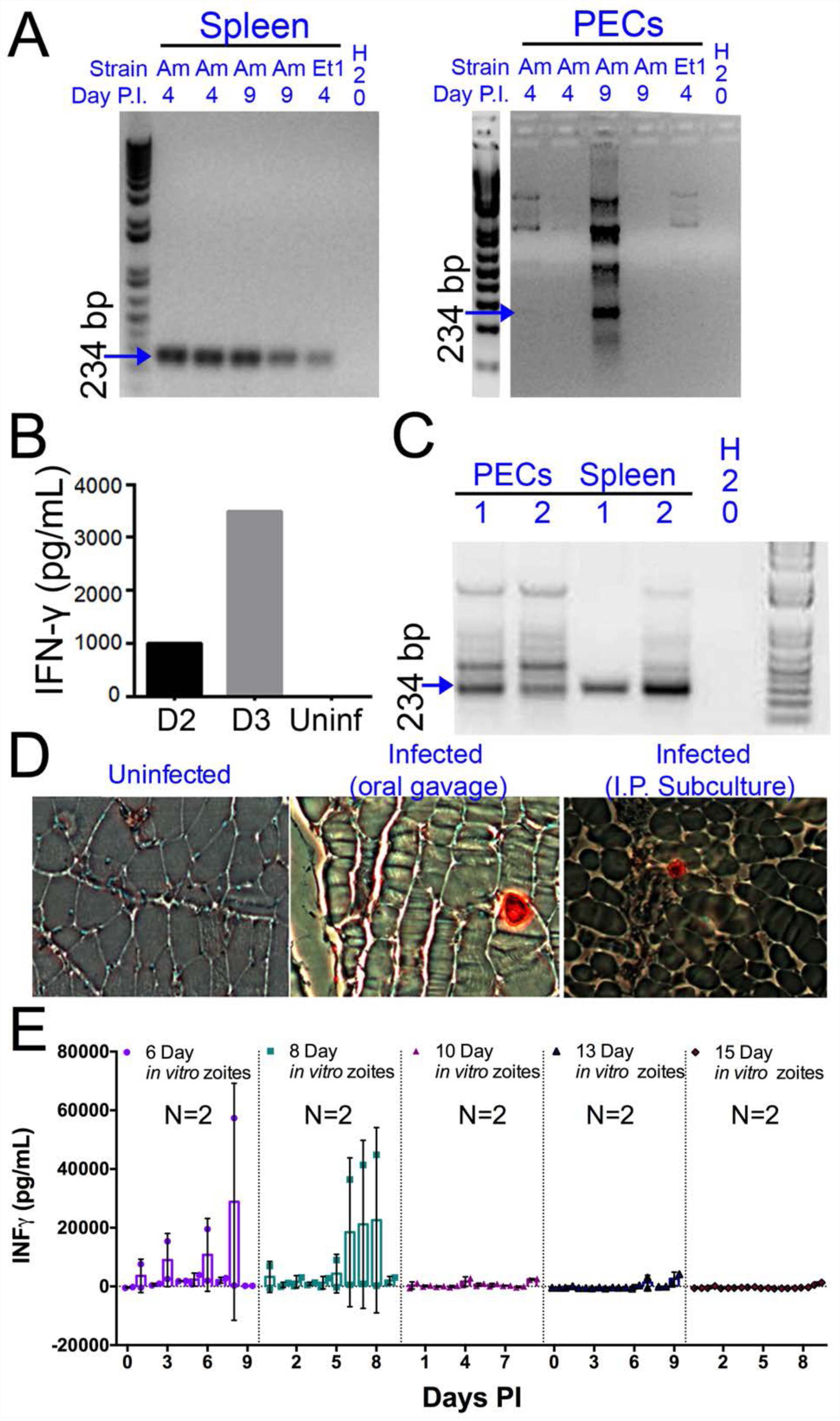
*H. hammondi* parasites can be grown *in vitro* and then used to successfully infect mice. A) Five Balb/c mice were infected intraperitoneally with *H. hammondi* zoites (HhCatAmer and HhCatEth1) grown in culture for 4 days. At either 4 or 9 DPI, parasite DNA was detected in the spleen and peritoneal lavage using PCR with *H. hammondi*-specific primers. Arrow indicates primary band for *H. hammondi*-specific PCR, although we also routinely see larger bands after PCR amplification. Samples were only considered positive if the 234 bp band was present (arrowhead). B-D) Two Balb/C mice were infected with ~50,000 *H. hammondi* zoites grown *in vitro* for 5 days. B) Serum was collected on days 2 and 3 and assayed for interferonγ. C) Three weeks post-infection, DNA was harvested from peritoneal lavage cells and spleen and assayed for *H. hammondi* DNA using *H. hammondi*-specific primers. D) Leg muscles from infected mice were sectioned and stained with *H. hammondi*-reactive anti *Toxoplasma* antibodies and compared to uninfected mice and mice infected with *H. hammondi* by oral gavage with 50,000 sporulated oocysts. E) *In vitro* cultivation leads to a dramatic and predictable loss in the ability to infect mice. Two BALB/C mice were infected with 50,000 *H.* hammondi American parasites after 6, 8, 10, 13, and 15 days of *in vitro* growth. The mass of the mice was monitored and a serum sample was obtained daily for 9 days post-infection. Analysis of interferon-γ production was analyzed in serum samples using an ELISA. Mice infected with parasites grown *in vitro* for 6 days (1 of 2) showed spikes in interferon-γ levels. This gamma spike was also observed in mice infected with parasites grown *in vitro* for 8 days (2 of 2). No gamma spike was observed in mice infected with parasite grown *in vitro* for 10, 13, or 15 days.

To determine if *in vitro* cultivated *H. hammondi* could initiate chronic infections in mice, we injected 5 Balb/c mice intraperitoneally with ~50,000 day 5 zoites. Serum collected on days 2 and 3 post-infection had detectable IFNγ, indicative of active infection (Fig 6B). At 3 weeks post-infection *H. hammondi* DNA was detected in both the spleen and peritoneal cell preparations from both mice (Fig 6C). Histological examination of muscle from infected mice using anti-*Toxoplasma* antibodies that cross-react with *H. hammondi* showed the presence of the parasites in the muscle tissue of the mouse infected with *in vitro*-passed *H. hammondi* (right panel, Fig 6D). These tissue-dwelling parasites were similar to those observed after oral inoculation with *H. hammondi* sporulated oocysts (middle panel; Fig 6D). Overall these data show that *H. hammondi* parasites grown *in vitro* are capable of a) inducing an immune response in mice during the acute phase of infection, and b) disseminating to multiple mouse tissues and establishing a chronic infection that resembles an infection resulting from oral gavage with sporulated oocysts (as shown in previous studies; (2, 15)).

After finding that *H. hammondi* zoites could infect mice after 4-5 days in culture, we aimed to determine if this ability changed over time. Mice were injected intraperitoneally with 50,000 zoites (released by needle passage) grown *in vitro* for 6, 8, 10, 13, and 15 days, and monitored for signs of infection by analyzing IFNγ levels in serum samples taken daily following infection. Remarkably, the ability to be subcultured *in vitro* tracked perfectly with the ability to initiate infections in mice *in vivo*. We observed an increase in IFNγ levels (indicative of infection) in 1 of 2 mice infected with 50,000 Day 6 zoites, and observed this spike in 2 of 2 mice infected with 50,000 Day 8 zoites. We did not see an increase in IFNγ levels in mice infected on with parasites cultured for 10, 13, and 15 days, suggesting that after 8 days of *in vitro* culture *H. hammondi* is no longer infectious to mice, at least to a degree that results in induction of detectable serum IFNγ levels (Fig 6E).

### *H. hammondi* parasites can be transfected using electroporation and grown *in vitro.*

Genetic manipulation of *H. hammondi* has not been reported in the literature. Given the genetic similarity between *T. gondii* and *H. hammondi* (11, 12), and the apparent interchangeability of promoter sequences between them (11, 23), we transfected ~4 million HhCatAmer sporozoites with a UPRT-targeting CRISPR/CAS9-GFP plasmid along with a PCR2.1 Topo-cloned *dsRED* cassette with 20 bp sequences flanking the CAS9 nuclease cut site within the *H. hammondi UPRT* gene. We used the *Toxoplasma*-specific plasmid to target *H. hammondi UPRT* as the gRNA sequence is identical between *T.gondii* and *H. hammondi*. As shown in Figure 7A, both of the plasmids were taken up and expressed by *H. hammondi*, as evidenced by GFP-fluorescence in the parasite nuclei and red fluorescence in the cytoplasm (Fig 7A). Importantly, transgenic parasites replicated *in vitro* as evidenced by the presence of multicellular vacuoles in the infected monolayer (Fig 7A). Transfection efficiency of viable parasites (as evidenced by those replicating *in vitro* with CAS9-GFP and/or dsRED staining) was ~4% (15 out of 350 vacuoles at 48 h post-transfection), which is consistent with typical transfection efficiencies observed with most strains of *T. gondii*. We reasoned that this system could be used to make a stable transgenic line of *H. hammondi*.

**Fig 7.**
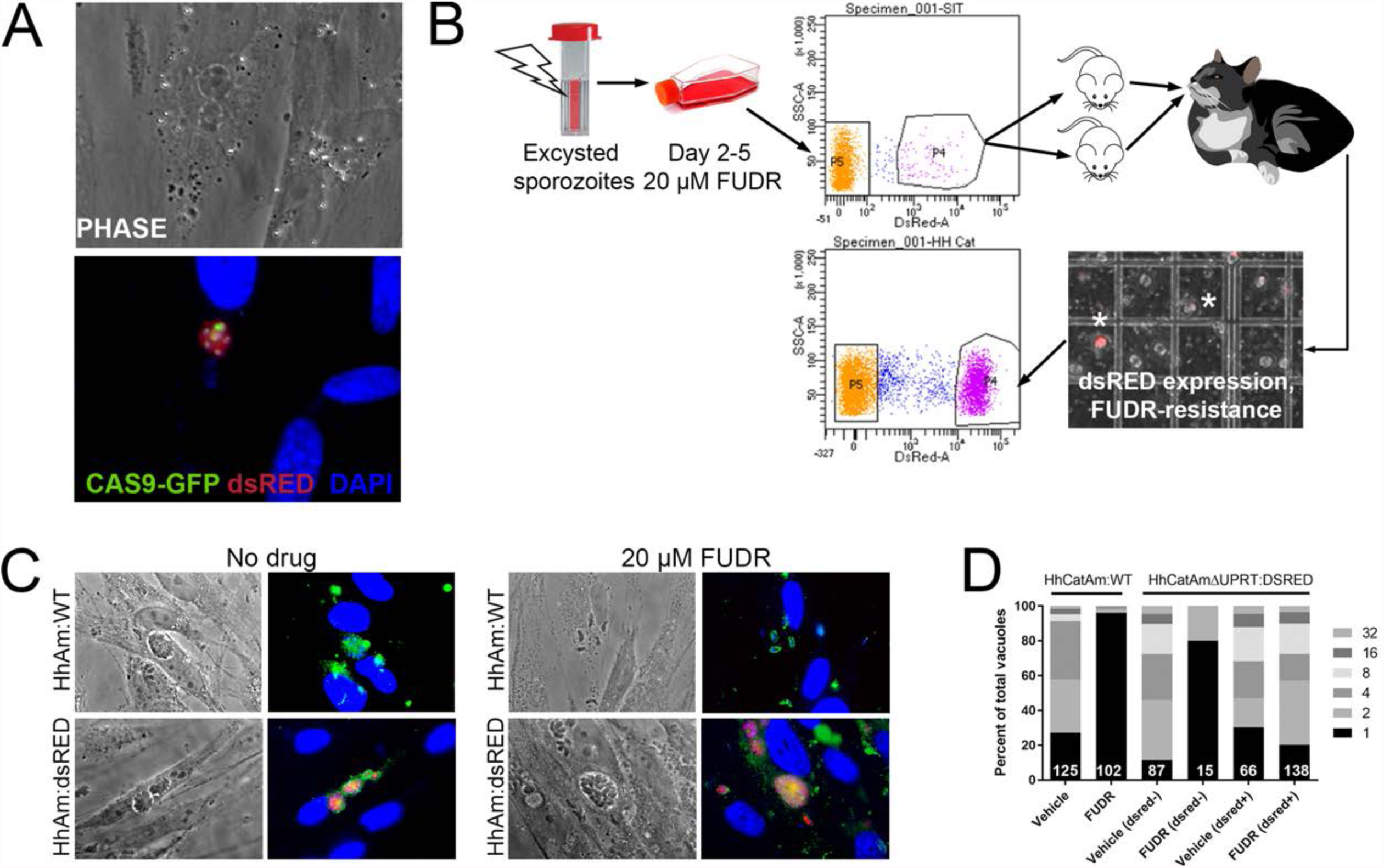
Generation of stable transgenic *Hammondia hammondi*. A) *H. hammondi* zoites co-transfected with CRISPR/CAS9-GFP and a plasmid harboring a *T. gondii* dsRED expression cassette. We identified multiple parasites with both GFP-tagged nuclei (due to CAS9-GFP expression) and dsRED-tagged cytoplasm, indicating uptake of both plasmids within the same parasites. B) Protocol for generating stably transgenic *H. hammondi* using flow cytometry and drug selection to enrich for transgenic parasites prior to cat infections. C,D) Testing FUDR resistance in transgenic *H. hammondi* parasites. WT and transgenic *H. hammondi* sporozoites were incubated in the presence or absence of 20 μM FUDR, and vacuole size and presence/absence of dsRED was quantified in at least 100 vacuoles per treatment condition. While WT and non-dsRED-expressing *H. hammondi* from the transgenic population were highly susceptible to FUDR treatment (as evidenced by the fact that the majority of vacuoles contained only 1 parasite), dsRED expressing transgenic parasites were resistant to FUDR treatment, confirming that the parasites harbored two genetic markers.

### Using *T. gondii* genetic tools to create a stable transgenic *H. hammondi* line

After transfection of ~4 million parasites with the Hh*uprt*:dsRED repair cassette and the gRNA*uprt*:CAS9:GFP plasmid, we cultured them in 10 μM FUDR during days 2-5 post-transfection. After FACS sorting for dsRED-expressing zoites, we sorted ~12,000 parasites (~3% of the total parasite population; Fig 7B), and injected two female Balb/c mice intraperitoneally with ~6000 dsRED-expressing zoites each. For the first week of the infection, mice were injected 1X daily with 200 μL of 1 mM FUDR, and both mice were euthanized and fed to a cat 4 weeks post-infection. Fecal floats showed the presence of dsRED-expressing sporozoites within some of the oocysts (Fig 7B), and ~32% of isolated sporozoites expressed dsRED based on FACS analysis (Fig 7B, bottom FACS plot). PCR screening (S2 Fig) and the inability to grow parasites long term *in vitro* confirmed the identity of the fluorescent parasites as *H. hammondia*.

To determine if our transgenic approach to disrupt the *UPRT* gene and insert a dsRED expression cassette was successful, we quantified the growth of dsRED^+^and dsRED^-^ parasites in the presence of FUDR and compared it to wild type HhCatAmer. Fixed and permeabilized parasites were stained with cross-reactive anti-*Toxoplasma* antibodies to identify all vacuoles. After 4 days of growth, we quantified vacuole size and dsRED-expression status in >100 vacuoles for each strain (WT and dsRED) and condition (Vehicle or 20 μM FUDR). As expected, HhCatAmer:WT grew poorly in FUDR (Fig 7C), while HhCatAmer:dsRED parasites grew similarly to untreated parasites (Fig 7C). When we separated parasite growth in FUDR based on dsRED expression, we found that dsRED^-^ parasites in the *H. hammondi* dsRED population were just as susceptible to FUDR as were HhCatAmer:WT parasites (Fig 7D, first 4 bars), while dsRED^+^ parasites grew equally well in FUDR compared to Vehicle (Fig 7D, right 2 bars). These data suggest that most, if not all, of the dsRED-expressing parasites are also null for the *UPRT* gene, either by direct replacement of the *UPRT* gene with the dsRED cassette or by incorrect non-homologous end joining and insertion of the dsRED cassette at another genomic location. This represents the first stably transgenic *H. hammondi* line.

### The *H. hammondi* transcriptome is enriched for bradyzoite-specific genes and genes expressed only in *T. gondii* cat enteroepithelial stages

Using existing genome annotations, we identified 7372 genes that were informatically determined to be syntenic orthologs between *T. gondii* and *H. hammondi* (ToxoDB.org). This is likely a conservative estimate of shared gene content between *T. gondii and H. hammondi* (12), but we used this dataset as a starting point for our transcriptional comparisons. The raw number of read counts mapping to coding sequences and overall transcript coverage from both the *T. gondii and H. hammondi* assemblies are shown in Figure 8A. As expected based on their dramatically different growth rates *in vitro* (Figs. 1 and 2), the number of cDNA reads mapping to *H. hammondi* coding sequences was lower overall compared to those mapping to *T. gondii*, and this was particularly pronounced at the 15 DPI. We addressed this issue by including genes in downstream analyses only if one *H. hammondi* D4 and one *H. hammondi* D15 sample had at least 5 reads. This reduced the overall size of the queried gene set to 4146, and all analyses were performed with this subset.

**Fig 8.**
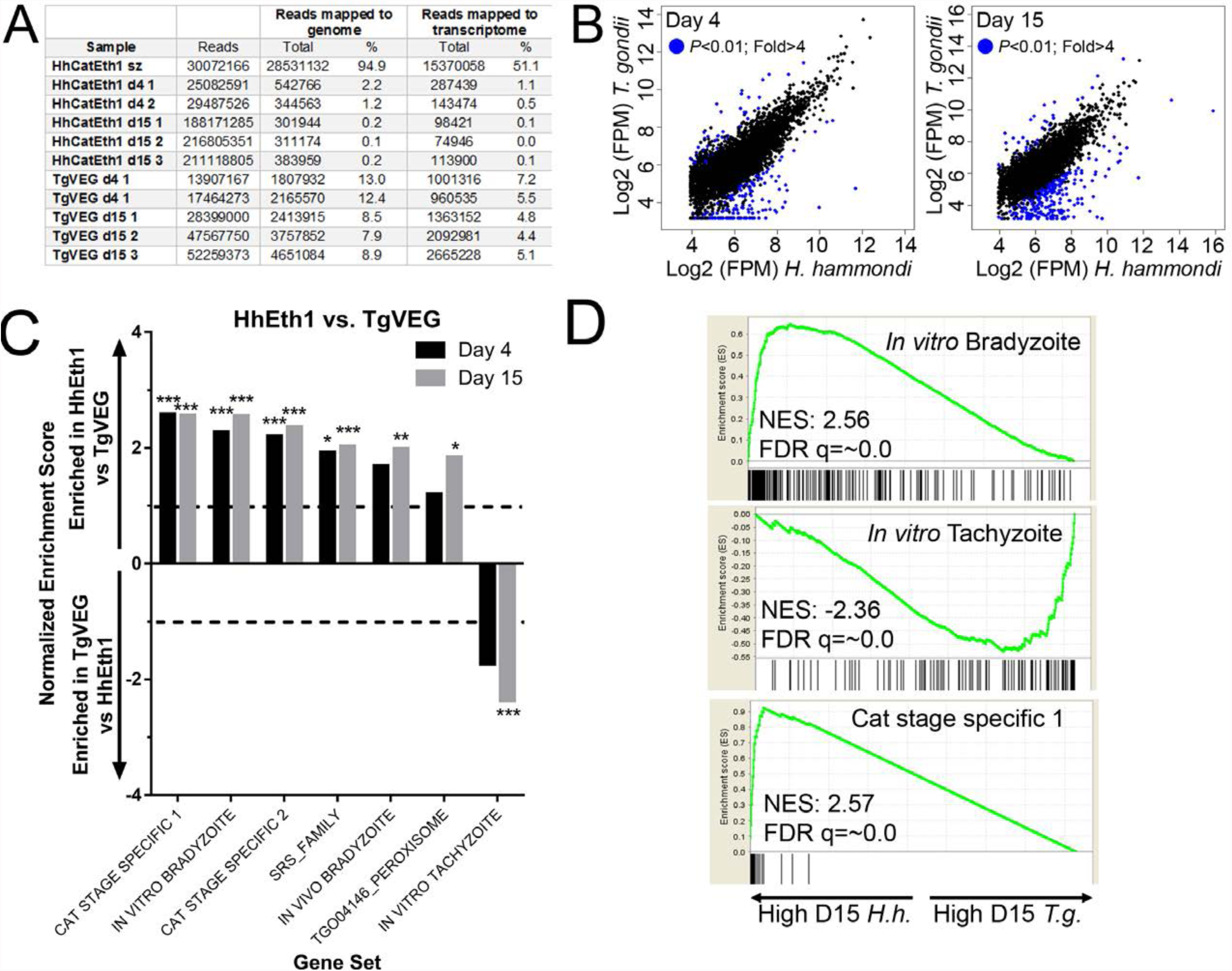
mRNAseq comparisons between *T. gondii* and *H. hammondi* identify unique aspects of the *H. hammondi* transcriptome. A) Summary of reads obtained for each parasite species and life stage, and the percentage of reads from each that mapped to the respective parasite genomes or transcriptomes. Log_2_-transformed and normalized Fragments Per Million (FPM) data for the 4276 shared genes between *T. gondii* and *H. hammondi* that passed thresholds for detection in both species. Genes of significantly different abundance (based on criteria as listed in inset) are blue. C) Gene Sets found to be significantly (FDR q-value < 0.05) enriched in *H. hammondi* high-abundance transcripts (top) or *T. gondii* high-abundance transcripts (bottom) at either 4 or 15 days post-excystation. Gene Set Details can be found in Table 2. *: P<0.05; **: P<0.01; ***: P<0.001. D) GSEA plots of the *In vitro* bradyzoite, *in vitro* tachyzoite, and Cat stage specific 1 gene sets, showing enrichment profiles between *H. hammondi* and *T. gondii* at 15 DPI. NES; Normalized enrichment score. FDR q: False discovery rate q-value.

**Table 2:**
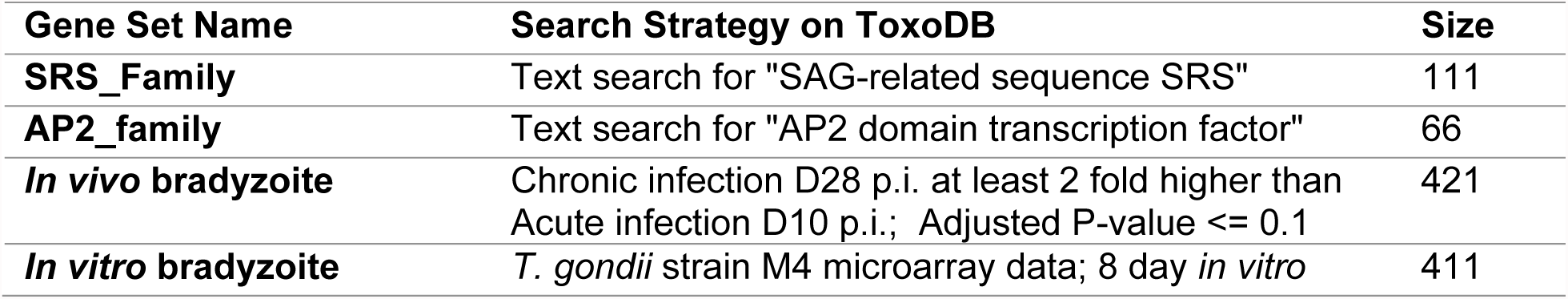
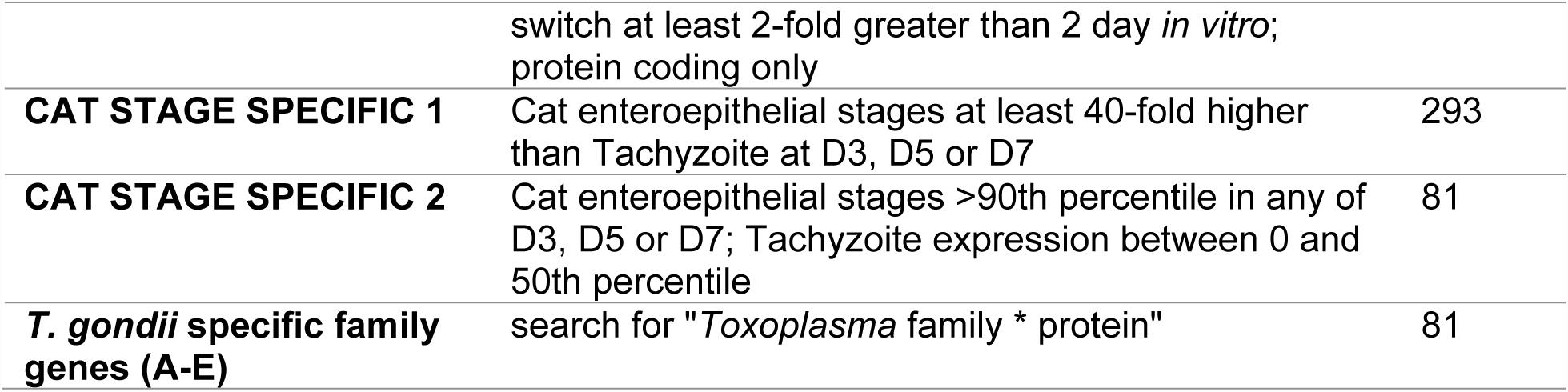
Supplemental gene sets used in Gene Set Enrichment analyses.

| Gene Set Name | Search Strategy on ToxoDB | Size |
| --- | --- | --- |
| **SRS_Family** | Text search for "SAG-related sequence SRS" | 111 |
| **AP2_family** | Text search for "AP2 domain transcription factor" | 66 |
| ***In vivo* bradyzoite** | Chronic infection D28 p.i. at least 2 fold higher than Acute infection D10 p.i.; Adjusted P-value $\leq 0.1$ | 421 |
| ***In vitro* bradyzoite** | *T. gondii* strain M4 microarray data; 8 day *in vitro* | 411 |
|  | switch at least 2-fold greater than 2 day *in vitro*;<br>protein coding only |  |
| **CAT STAGE SPECIFIC 1** | Cat enteroepithelial stages at least 40-fold higher<br>than Tachyzoite at D3, D5 or D7 | 293 |
| **CAT STAGE SPECIFIC 2** | Cat enteroepithelial stages >90th percentile in any of<br>D3, D5 or D7; Tachyzoite expression between 0 and<br>50th percentile | 81 |
| ***T. gondii* specific family**<br>**genes (A-E)** | search for "*Toxoplasma* family * protein" | 81 |

When log_2_ transformed and DESeq2-normalized RPM (mapping Reads Per Million; all data in Table S1) values were plotted for Day 4 and Day 15 between species we observed a high correlation in transcript levels at both time points (Fig 8B), similar to our previous results with sporulated oocysts (23), with a clear subset of genes with differential expression (blue data points, Fig 8B). The majority of genes were expressed at similar levels between *T. gondii* and *H. hammondi* (hierarchically clustered heat map shown in S3A Fig), although subclusters of transcripts of different abundance are clearly present (annotated with lines and labels in S3A Fig). We used Gene Set Enrichment Analysis (GSEA; (24)), a set of previously published (25) gene sets, and supplemented them with our own gene sets to compare Day 4 and Day 15 *T. gondii* and *H. hammondi* transcriptional profiles (see Methods and Table 2 for Gene Sets).

When comparing *T. gondii* and *H. hammondi* at each time point, we identified 7 gene sets that showed significant enrichment among *H. hammondi*-high or *T. gondii*-high genes (Fig 8C). For *H. hammondi*, profiles showed significant enrichment for *in vitro* (D4, D15) and *in vivo* (D15) bradyzoite gene sets (Fig 8C,D), among others. This is consistent with the spontaneous bradyzoite differentiation phenotype characteristic of *H. hammondi* described above. Consistent with *H. hammondi* spontaneous differentiation, *T. gondii* showed significant enrichment for the *in vitro* tachyzoite gene set at day 15 p.i. (Fig 8C,D), a time point at which *H. hammondi* is over 50% dolichos-positive (Fig 3). We also assayed for gene sets that changed within each species between the D4 and D15 time points (S3B Fig). For the most part, *T. gondii* and *H. hammondi* shared a similar gene set enrichment profile at the D15 time point, sharing an enrichment for *in vitro*/*in vivo* bradyzoite genes and the SRS_family compared to D4 (S3B Fig).

A more unexpected finding was that at both day 4 and 15, *H. hammondi* had a transcriptional profile with highly significant enrichment for genes that are uniquely expressed in *T. gondii* cat enteroepithelial cells ((26, 27); e.g., merozoite-specific genes; CAT STAGE SPECIFIC 1, 2; Fig 8C,D)). Specifically, out of 21 members of the “CAT STAGE SPECIFIC 1” gene set that were detectable, 18 of them were ranked in the top 308/4146 (7.4%) and 174/4146 (4.2%) of *H. hammondi*-high genes at D4 and D15, respectively (Fig 8D). These data suggest that *H. hammondi* has aspects to its transcriptional profile that are independent of its spontaneous conversion to bradyzoites and that a subset of genes originally thought to be restricted to merozoites may only be merozoite-specific in some species and not others.

Using DESeq2, we identified 344 genes of significantly different abundance between species at either the D4 or D15 time points, and 126 of these were found to be significant at both time points. We then focused on the enriched gene sets identified by GSEA (Fig 8C,D) and assessed the degree of overlap at each time point for significantly regulated genes from the *in vitro* bradyzoite, *in vitro* tachyzoite, and merozoite-specific gene sets (Fig 9A). While bradyzoite and *T. gondii* merozoite-specific transcripts were significantly higher in *H. hammondi* at D4 and D15 (Fig 9A-B,D), the bulk of the tachyzoite-specific genes found to be significantly higher in *T. gondii* compared to *H. hammondi* were restricted to the D15 time point (Fig 9A,C). This is consistent with the robust conversion to dolichos-positive bradyzoites in *H. hammondi* compared to the tachyzoite-like proliferation of *T. gondii* VEG.

**Fig 9.**
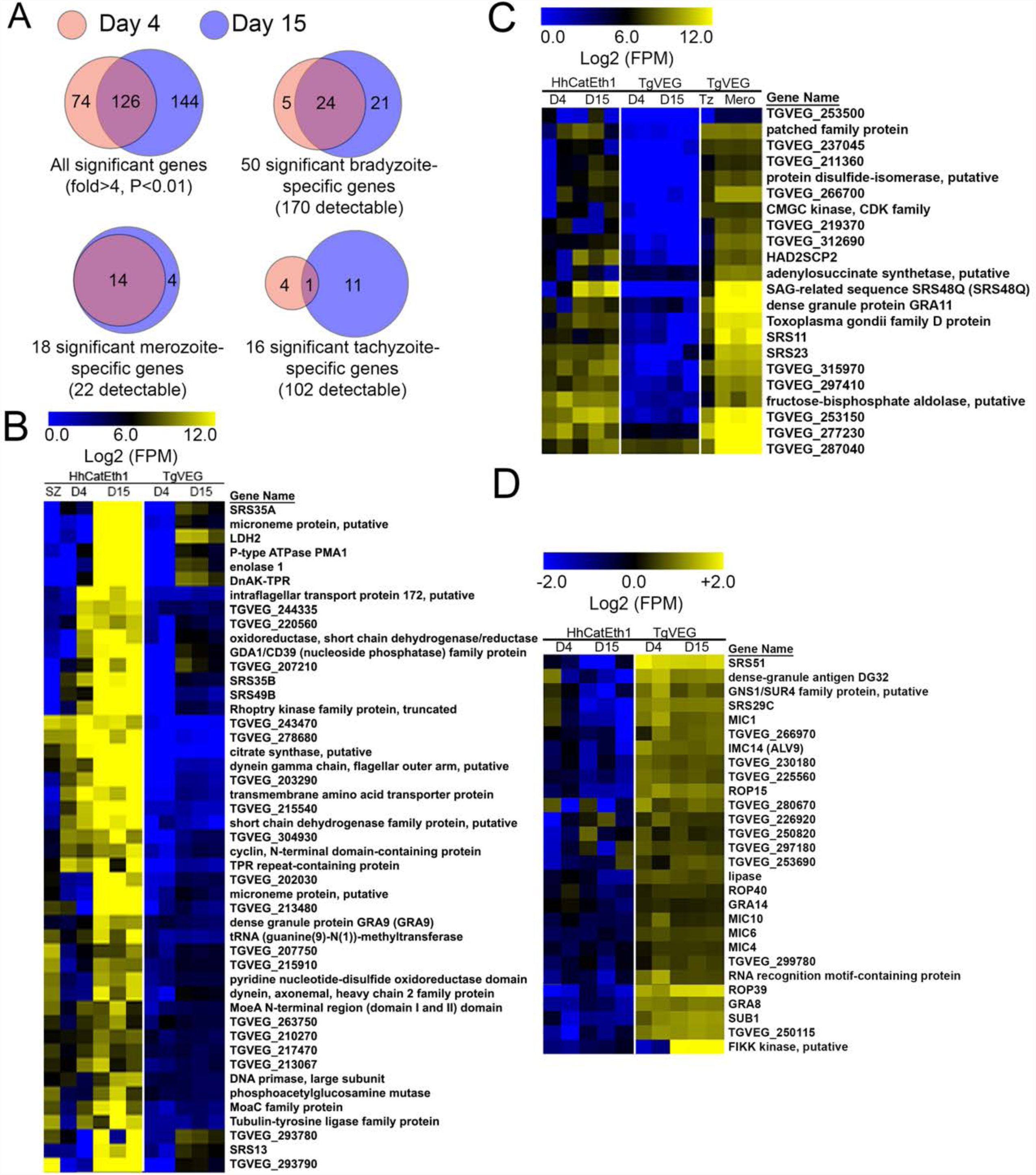
Analysis of the overlap between genes found to be of significantly different abundance at D4 and D15 PI across different gene sets. A) While overall there are distinct genes that are significantly different at D4 and D15 (Venn diagram, upper left), *H. hammondi* expresses high numbers of bradyzoite genes even at 4 dPI and this number increases further by D15 (21 additional genes; Venn diagram in upper right). This progression towards an increase in the number of bradyzoite genes over time is not recapitulated among merozoite-specific genes, in that only 4 additional detectable genes from this gene set were found to be of significantly higher abundance at 15 DPI. Moreover 18 of the 22 total detectable merozoite genes were of significantly different abundance in *H. hammondi*, indicating its transcriptional similarity to both bradyzoites and merozoites. B-D) Heat maps showing detectable genes from the bradyzoite, tachyzoite and merozoite-specific gene sets. B) Bradyzoite genes (raw log_2_ FPM data shown). C) Merozoite-specific genes (raw Log_2_ FPM data shown). D) Tachyzoite-specific genes (mean centered Log_2_ FPM data shown).

One caveat with these data is that the number of reads mapping to the *H. hammondi* transcriptome was, on average, an order of magnitude lower than those that mapped to *T. gondii*. While this is most certainly due to the dramatic differences in replication rate between these species (Fig 2), we validated a subset of transcripts that were of greater abundance in *H. hammondi* using qPCR. To do this, we used *ROP18* as a positive control for a gene that was of significantly higher abundance in *H. hammondi* compared to *T. gondii* (TgVEG and other Type III strains express very little *ROP18* transcript; (11, 23, 28)), and *CDPK1* as a negative control for a gene of similar abundance between the two species (S4A Fig). *GRA1* transcript levels were used as loading controls. After validating primer sets (which had to be made separately for each species; see Table S2 for all primer sequences), we quantified 8 transcripts (plus *ROP18*) using qPCR on RNA freshly isolated from D4 and D15 HhCatEth1 and TgVEG cultures in HFFs. As expected *ROP18* was found to be >1000-fold greater in abundance in HhCatEth1 D4 samples and >16-fold higher in D15 samples, while *CDPK1* transcript level was not significantly different between species at either time point (S4B Fig). The remaining 8 queried transcripts showed significantly higher abundance in *H. hammondi* compared to *T. gondii* at D4, and 2 were found to also be of significantly higher abundance at the D15 time point (S4B Fig). Therefore, we overall confirmed the increased transcription of a subset of queried genes between *T. gondii* and *H. hammondi* during *in vitro* culture, suggesting that our approach to identify transcripts of significantly higher abundance had a relatively low Type I error rate, despite the fact that there was such a large difference in the number of reads mapping to the *H. hammondi* transcriptome compared to *T. gondii* (Fig 8A).

## Discussion

The ability to infect multiple hosts is a remarkable feature of many parasite species. Parasites with multi-host life cycles undergo stereotyped patterns of development and growth in a given host, reaching a unique developmental state in the life cycle that is transmissible to the next host. With very few exceptions, parasites with heteroxenous life cycles are obligately so; that is to say that life stages infectious to one host (e.g., the definitive host) are not infectious to another host (e.g., an intermediate host). Among the Sarcocystidae (including such parasites as *Sarcocystis spp.*, *Besnoitia spp.*, *Neospora spp.*), all have exogenous life cycles for which they are obligately heteroxenous with the exception of *T. gondii*. Only in *T. gondii* can oocysts and tissue cysts (i.e., bradyzoites) infect, and ultimately be transmitted by, both intermediate and definitive hosts. In addition to its comparatively flexible life cycle, *T. gondii* is more virulent in the mouse model than *H. hammondi* (for example, IFNγ-knockout mice survive infection with *H. hammondi* while *T. gondii* is highly lethal in this host genetic background (2, 29)).

Given that the molecular determinants of life cycle and host range are poorly understood in most parasite species, we are exploiting the *T. gondii*/*H. hammondi* system as a way of understanding how complex life cycles evolve, are maintained, and can be altered. In the present study, we conducted comparisons between one strain of *T. gondii* (VEG strain) and 2 strains of *H. hammondi* (HhCatEth1 and HhAmer), quantifying differences in excystation frequency, infectivity *in vitro* and *in vivo*, replication rate, and development. Ideally, one *H. hammondi* strain would be used for all experiments, but two strains of *H. hammondi* were utilized due to oocyst availability at the time of experimentation. We identified dramatic differences in parasite biology between these two species that likely contribute to their differences in pathogenesis in the mouse model, and that may also be important in determining the host species in which this parasite can cause disease.

The nearly 2-fold difference in replication rate could play an important role in parasite virulence (Table 1, Fig 2D, E), assuming this phenotype is recapitulated *in vivo*. Besides the impact that a slower replication rate would have on acute virulence, it may also impact infection outcome in the chronic phase. While *H. hammondi* parasites can at times be detected in the brains of mice after infection with oocysts (2), this particular outcome that is a hallmark of *T. gondii* infection (2, 30) is much less common after exposure to *H. hammondi*. It is possible that *H. hammondi* is unable to cross the blood-brain barrier via infection of endothelial cells (31) or is poorly trafficked by dendritic cells and/or macrophages. However, a simple explanation for this difference in “tropism” could be driven solely by the comparatively lower parasite burden during *H. hammondi* infection in mice. In contrast to differences in replication rate, we observed no substantial differences in the ability of *T. gondii* and *H. hammondi* to invade and form vacuoles within HFFs, suggesting that their invasion machinery is mostly conserved and intact. This idea finds some support in our transcriptome data, where we detected no enrichment for gene sets consisting of microneme, rhoptry or dense granule genes, and detected similar amounts of crucial invasion genes like AMA1, SpAMA1, as well as multiple rhoptry neck proteins (see Table S1 for normalized RNAseq data).

One surprising finding was that we repeatedly harvested significantly higher sporozoite yields from *T. gondii* compared to *H. hammondi* (~2-fold overall; Fig 1A), and this was independent of oocyst age and the preparation of *H. hammondi* (Fig 1B). One caveat is that the same prep of TgVEG oocysts was used for all 10 extractions, so this observation could be due to differences in this parasite preparation. However, we recently excysted a separate, 4 month old TgVEG oocyst preparation and obtained a ~13% sporozoite yield, suggesting that the ability to excyst may be a consistent difference between the species. While the excystation procedure is artificial, if *T. gondii* excysts more readily than *H. hammondi,* this could lead to higher infection efficiencies and ultimately more robust transmission. The comparative ease of excystation could also be related to the ability of *T. gondii* oocysts to excyst in both definitive and intermediate hosts. This might compromise environmental stability (due to a less robust oocyst or sporocyst wall), but the trade-off with respect to increased host range and transmission rates might be immense and highly favored. Experiments examining excystation rates within the animal itself and environmental stability will be necessary to address this question directly.

While others (15) have observed that *H. hammondi* grown in tissue culture for many weeks transformed into infectious cyst stages, the exact timing of this event was not known. Spontaneous cyst conversion after initiating cultures with *T. gondii* sporozoites have also been described, and in most detail for *T. gondii* strain VEG (13). In contrast to TgVEG, which showed varying rates of spontaneous conversion but continued to replicate throughout the experiment, *H. hammondi* (American strain) fully converted to a bradyzoite cyst-like stage by 23 days post infection (DPI). This programmed differentiation correlated with the ability of *H. hammondi* to infect new host cells *in vitro* and i*n vivo.* Our results suggest that *H. hammondi* is capable of subculture for a limited, yet highly predictable window. This finding is in contrast to previous studies where subculture was unsuccessful (14, 15). However, in these studies, subculture of *H. hammondi* was attempted after 1, 2, 4, and 6 weeks of growth, and all but one of these time points are outside of the period which we described as the window of infectivity for *H. hammondi*. Furthermore, insufficient details regarding experimental approach to subculture and infection analysis could make it possible that infection occurred and was undetected, or that not enough parasites were used for subculture to allow for detection of parasites. This explanation is supported by the dramatic decline in the number of observable parasites near the end of our 8 day window of subculture for *H. hammondi.* When taken together with the dynamics of spontaneous differentiation in *H. hammondi*, these data demonstrate that *H. hammondi* terminally differentiates into a life stage that is no longer infectious to anything but the definitive host, including neighboring host cells. It’s unknown whether terminally differentiated *H. hammondi* tissue cysts could be coaxed to infect cat gut epithelial cells *in vitro.* This would be a novel approach to determine the relative importance of host cells versus the host cell environment in determining life cycle restrictions. *T. gondii* does not appear to have a life stage that is analogous to the *H. hammondi* terminally differentiated tissue cyst. Both *in vivo-* and *in vitro*-generated *T. gondii* tissue cysts can recrudesce to the tachyzoite stage, leading to lethal disease in immunocompromised and organ transplant patients (32, 33). While it is not known if *H. hammondi* can infect humans, based on the data in hand, we predict that if human infections with *H. hammondi* did occur, tissue cysts would be unable to cause disease in immunosuppressed or immunocompromised patients due to a) low or non-existent recrudescence rates and/or b) inability of recrudesced parasites to invade neighboring host cells. Further work in mouse reactivation models could help to address this question, in addition to the development of serological tests that could immunologically distinguish these two parasites.

*T. gondii* of nearly all genetic backgrounds can be induced to form tissue cysts *in vitro* through the application of a variety of stressors (pH, nitric oxide donors, serum starvation; (13, 34, 35)), and conversion to tissue cysts in *T. gondii* facilitates effective transmission since the cyst wall aids in survival of the parasite as it passes through the stomach. Given the importance of the stress response in *T. gondii* replication and transmission dynamics, we were surprised to find that day 2 *H. hammondi* zoites exposed to high pH (8.2) medium for 2 days did not form any DBA-positive cysts. This suggests that *H. hammondi* may less sensitive to external stressors than *T. gondii*, despite the fact that it ultimately converts to a 100% DBA-positive population. It is possible that being responsive to external stress facilitates a flexible *T. gondii* life cycle, enabling it to increase its population to the highest possible level (depending on the immune response of the host) prior to converting to the tissue cyst stage. In the case of *H. hammondi*, the lack of stress-induced switching may reflect that *H. hammondi* is predictably moving towards terminal differentiation, and therefore is under no selective pressure to sense the host environment (or at least no pressure to respond to those changes by converting to a cyst). If the progression from replicating *H. hammondi* zoite to tissue cyst is as predictable *in vivo* as it is *in vitro*, timing is crucial for this strategy to be successful since it is thought that only bradyzoites are immune to the toxic host response. It may be that the timing of *H. hammondi* conversion is compatible with rodent immune responses (known intermediate hosts for *H. hammondi*; (14, 22, 36)), but may not be in other species. The *T. gondii*/*H. hammondi* comparative system provides a unique opportunity to address these questions more directly.

Some insight into the *H. hammondi* developmental process can be gleaned from the 4 and 15 DPI transcriptional profiling data. *H. hammondi* consistently upregulated genes previously defined as being uniquely expressed in *T. gondii* bradyzoites (whether *in vivo* or *in vitro*) as early as 4 DPI, indicating that there is a high degree of similarity between the spontaneous differentiation process that occurs in *H. hammondi* and the stress-induced process observed in *T. gondii* (25, 37, 38). This observation is consistent with the emergence of DBA-positive *H. hammondi* vacuoles throughout the developmental process *in vitro*. Less expected, however, was the constitutive expression of genes in *H. hammondi* that are restricted to cat enteroepithelial stages in *T. gondii* (27). It is exciting to speculate that the expression of certain genes in *H. hammondi* bradyzoites that are merozoite-specific in *T. gondii* is related to the fact that *H. hammondi* bradyzoites are only infectious to cat cells. Future studies aimed at obtaining higher coverage transcriptomes will help to address this, as will the use of our newly developed transgenic system for *H. hammondi*.

In contrast to previously published work, we identified a short but very predictable replicative window during which *H. hammondi* could be used to infect new host cells (whether those cells were in an animal or a tissue culture dish). We exploited this time period to genetically manipulate *H. hammondi* using gene editing strategies developed for *T. gondii.* (39) Using these strategies, along with our knowledge of the limits of *H. hammondi’s in vitro* growth, we generated the first transgenic *H. hammondi* parasites expressing dsRED (30-40% after the first round of selection) that were >99% resistant to FUDR. This work represents an important step in improving the genetic tractability of the *H. hammondi* system, paving the way for future studies targeting specific loci that may be implicated in the dramatic differences in the life cycles, lytic cycles, and virulence determinants of these closely-related parasite species.

## Materials and methods

### Parasite strains and oocyst isolation

Oocysts of *Hammondia hammondi* strain HhCatEth1 (18), HhCatAmer (22) and *Toxoplasma gondii* strain VEG (16) were isolated from experimentally infected cats as described previously (16). Briefly, wild type (for *T. gondii*) or interferon-γ (IFN-γ) knockout (for *H. hammondi*) mice were orally infected with 10^4^ sporulated oocysts and sacrificed 4-6 weeks post-infection, and leg muscle (for *H. hammondi*) or brain (*T. gondii*) tissue were fed to 10-20 week old specific pathogen-free cats. Feces were collected during days 7-11 post-infection, and unsporulated oocysts were isolated via sucrose flotation. Oocysts were placed at 4°C in 2% H_2_SO_4_ to allow for sporulation to occur and for long-term storage.

Due to the limited availability and difficulty associated with the generation of oocysts, two strains (HhCatEth1 and HhCatAmer) of *H. hammondi* generated from multiple cat-derived oocysts preparations were used in experiments described below. Furthermore, various MOIs were utilized throughout the experiments described below to compensate for differences in replication between *T. gondii* and *H. hammondi.*

### Excystation of T. gondii and H. hammondi Oocysts

Sporulated oocysts were washed 3X in Hanks’ balanced salt solution (HBSS; Life Technologies, 14175145) and treated with 10% bleach in PBS for 30 minutes while shaking at room temperature. Washed pellets were resuspended in 3 mL of HBSS in a 15-mL falcon tube, and 4g sterile glass beads (710-1,180 uM; Sigma-Aldrich, G1152) were added. Parasites were vortexed on high speed for 15 seconds on/15 seconds off, 4X. Supernatant was removed and pelleted by centrifugation. The pellet was resuspended and syringe-lysed using a 25 gauge needle in 5 mLs of pre-warmed (37°C) and freshly made, sterile-filtered excystation buffer (40 mL PBS, 0.1 g Porcine Trypsin (Sigma-Aldrich, T4799), 2 g Taurocholic Acid (Sigma-Aldrich, T4009), pH 7.5). After 45 minutes in a 37°C water bath, the suspension was syringe lysed again, and 7 mLs of cDMEM (100U/mL penicillin, 100μg/mL streptomycin, 2mM L-glutamine, 10% FBS, 3.7g NaH_2_CO_3_/L, pH 7.2) was added to quench the excystation media. Mixture was centrifuged, pellet resuspended in cDMEM, and passed onto monolayers of human foreskin fibroblasts (HFFs) grown at 37°C in 5% CO_2_.

### Fixing parasites for immunofluorescence assays

Parasites were washed twice with PBS, fixed with 4% paraformaldehyde in PBS (Affymetrix, 19943), washed twice with PBS, and blocked/permeabilized in blocking buffer (50mL PBS, 5% BSA, 0.1% Triton X-100).

### Quantification of sporozoite viability and replication rate

After 10 minutes incubation on HFFs at 37°C in 5% CO_2_, monolayers containing freshly excysted parasites were scraped, serially syringe lysed (25 and 27 gauge needles), and pelleted. The pelleted parasites were resuspended in cDMEM, filtered through a 5 μM syringe-driven filter (Fisher Scientific, SLSV025LS) and used to infect 4-chambered slides (Lab-Tek^®^ II, 154526) containing a confluent monolayer of HFFs at an MOI of 0.5 for both *H. hammondi* CatEth1 and *T. gondii* VEG. After 1, 2, and 3 days of growth, cells and parasites were fixed as described above and stored in blocking buffer at 4°C until needed. Cells were immunostained with goat polyclonal *Toxoplasma gondii* Antibody (ThermoFisher Scientific, PA1-7256) at 1:500. The secondary antibody, Alexa Fluor^®^ 594 Donkey anti-Goat IgG (H+L) (ThermoFisher Scientific, A-11037) was used at 1:1000. Coverslips were mounted to 4-chambered slides using VECTASHIELD Antifade Mounting Medium with or without DAPI, depending on the application (Vector labs, H-1000).

Vacuole formation was used as a proxy for sporozoite invasive capacity. To quantify this, the number of vacuoles with at least 1 parasite was determined in 100 random FOVs (1000x magnification) for 3 technical replicates for each parasite species at 1, 2, and 3 DPI. The entire experiment was repeated twice with different cat-derived oocyst preparations. For replication rate quantification, images of parasite-containing vacuoles were taken with an Axiovert 100 inverted fluorescent microscope with Zen lite 2012 software. The number of parasites per vacuole was determined for each image (biological replicates 1 & 2), and vacuole size (Biological replicate 1 only) was determined for each image using ImageJ software (NIH).

### Quantification of spontaneous *Dolichos biflorus-*positive cyst formation *in vitro*

For *Dolichos Biflorus* Agglutinin (DBA) staining, monolayers containing freshly excysted parasites were scraped, syringe lysed, and pelleted 24 hours after growth. The pellet was resuspended in cDMEM, filtered through a 5 μM syringe-driven filter (Fisher Scientific, SLSV025LS), and passed at MOIs of 0.3 (HhCatEth1) and 0.001 for (TgVEG) onto coverslips containing a confluent monolayer of HFFs. Three coverslips were infected per strain for each time point analyzed. At 4 and 15 DPI, cells and parasites were fixed as described above and stored in blocking buffer at 4°C until immunostaining was conducted.

In addition to being stained with rabbit *Toxoplasma gondii* Polyclonal Antibody (Invitrogen, PA1-7252) at a dilution of 1:500 and a 1:1000 dilution of Alexa Fluor^®^ 594 goat anti-rabbit IgG (H+L) (ThermoFisher Scientific, A-11037), coverslips were stained with Fluorescein labeled *Dolichos biflorus* Agglutinin (DBA; Vector Labs, FL-1031) at a dilution of 1:250. Coverslips were then mounted to microscope slides using ProLong^®^ Diamond Antifade Mountant with DAPI (ThermoFisher Scientific, P36962).

The number of DBA positive vacuoles was quantified in 20 FOVs in 3 coverslips for each parasite at 4 and 15 DPI. Images were obtained with an Axiovert 100 inverted fluorescent microscope with Zen lite 2012 software and edited using ImageJ software (NIH).

### Induction of bradyzoite formation in *T. gondii* and *H. hammondi*

Monolayers containing freshly excysted parasites were scraped, syringe lysed, and filtered through a 5 μM syringe-driven filter (Fisher Scientific, SLSV025LS) after 24 hours of growth. Filtered parasites were pelleted and passed at MOIs of 0.5 (*H. hammondi*) and 0.1 for (*T. gondii*) onto coverslips containing a monolayer of HFFs. Three coverslips were infected per parasite per treatment group. After 2 days, media was changed to pH 8.2 bradyzoite switch media (DMEM with 100U/mL penicillin, 100μg/mL streptomycin, 2mM L-glutamine 10mM HEPES, 2g/L NaHCO_3_, and 1% FBS (40)) or cDMEM. Coverslips with pH 8.2 media were grown at 37°C in the absence of CO_2_. Media was changed again at 3 DPI. After 4 DPI, cells and parasites were fixed as described above and stored in blocking buffer at 4°C until needed. Dolichos and counter-staining was conducted using Fluorescein labeled DBA and rabbit *Toxoplasma gondii* Polyclonal Antibody as described above. The number of DBA-positive vacuoles was quantified in 15 FOVs in 3 coverslips for each strain grown at either pH 7.2 or pH 8.2 and the percentage of DBA positive vacuoles was determined. Images were obtained and analyzed as described above.

### Subculture of *H. hammondi*

Sporozoites were obtained using the excystation protocol described above with the exception of syringe lysis and previously described host cell incubation. Six confluent monolayers of HFFs grown in T-25’s were infected with *H. hammondi* American (HhCatAmer) sporozoites at an MOI of ~2 (2.8 million sporozoites). After a three-hour incubation at 37°C, the Day 0 Transfer flask was scraped, syringed lysed 5X with 25 gauge needle, filtered through a 5 μM syringe-driven filter (Fisher Scientific, SLSV025LS), and passed to a new confluent monolayer of HFFs grown in a T-25. This process was repeated after 2, 5, 8, 11, and 15 days post-excystation. After 3 days of growth following passage, 4.17cm^2^ of the T-25 was monitored daily. Images of vacuoles were obtained with an Axiovert 100 inverted fluorescent microscope with Zen lite 2012 software with a 40X objective, and vacuole sizes were quantified using ImageJ software (NIH).

### Characterizing limits of *in vivo* infectivity of *H. hammondi* grown *in vitro*

Sporozoites were obtained using the excystation protocol described above. After 24 hours, HFF monolayers infected with excysted sporozoites were scraped, syringe lysed 5X with a 25 gauge needle, and pelleted. The pellet was resuspended in cDMEM, filtered through a 5μM syringe-driven filter (Fisher Scientific, SLSV025LS) and passed onto confluent monolayers of HFFs grown in T-25s. Infected host cells were incubated at 37 °C 5% CO_2_ from 6, 8, 10, 13, and 15 days. The media was replenished after 5-7 days. At each time point, infected host cells were scraped, syringe lysed 3X with a 25 gauge needle and 3X with a 27 gauge needle, and pelleted. Parasites were counted and diluted in PBS, and a dose of 50,000 parasites was injected intraperitoneally in 2 BALB/C mice for each time point. Serum samples were obtained from infected mice daily for 9 days post-infection and the mass of the mice was also monitored daily. After 9 days of infection, the mice were sacrificed and dissected. Tissue samples were preserved in 10% neutral buffered formalin (Sigma HT501128) until immunohistochemistry analysis was performed.

### Tissue sectioning and staining

Fixed tissue samples were embedded in paraffin, sectioned, and stained using rabbit anti-*Toxoplasma* antibody (ThermoScientific Cat# RB-9423-R7) by Research Histology Services at the University of Pittsburgh. For antigen retrieval, deparaffinized slides were steamed at pH=6.0 in 10 mM Citrate buffer. After exposure to 3% H_2_O_2_, slides were washed in Tris-buffered saline with 2.5% Tween-20 (TBST), and blocked for 20 min each with Avidin and Biotin blocking reagents (Vector labs; SP-2001) with TBST washes in between. Slides were blocked in 0.25% casein PBS for 15 min, and incubated overnight in primary antibody (1:100 dilution in 3% goat serum in PBS). Slides were washed in PBST and incubated for 30 minutes with biotinylated goat anti-rabbit (Vector laboratories; BA-1000; 1:200 dilution in 3% goat serum in PBS). Following 3 washes with PBST, slides were incubated for 30 minutes with streptavidin-HRP (Vector laboratories; PK-6100), washed 3X with PBST, and incubated with AEC substrate (Skytec; ACE-500/ACD-015) for 15 minutes. Following rinses in water, slides were counterstained with aqueous hematoxylin and blued using Scott’s tapwater substitute. Slides were mounted in Crystal Mount.

### Interferon-gamma (IFN-γ) ELISA

Blood samples were obtained from mice daily via submandibular bleed, allowed to clot, and centrifuged at 100xg. Serum was stored at -20°C until ELISAs were performed. IFN-γ levels were determined using the BD OptEIA™ Set Mouse IFN-γ kit (Cat.# 555138) according to manufacturer’s instructions. Serum samples were typically diluted 1:20.

### Transfection of *Hammondia* parasites and selection of recombinant parasites

Excysted sporozoites were prepared as described above and incubated overnight in a T25 flask with confluent monolayer of HFFs. After 24 hours, the monolayer was scraped, syringe lysed 3X with a 25 gauge and 27 gauge needle, and filtered using a 5 μM syringe-driven filter (Fisher Scientific, SLSV025LS). The filtered contents were pelleted by centrifugation at 800 x g for 10 min, and the pellet was resuspended in 450 μl of cytomix with 2 mM ATP and 5 mM glutathione. Resuspended parasites were transferred to a cuvette, electroporated at 1.6 KV and a capacitance of 25 μF, and used to infect confluent HFF monolayers on coverslips. The coverslips were fixed 5 DPI as described above and mounted using ProLong^®^ Diamond Antifade Mountant with DAPI (ThermoFisher Scientific, P36962).

To create stable transgenic *H. hammondi* parasite lines, the CRISPR/CAS9 plasmid expressing a gRNA sequence targeting the uracil phosphoribosyl transferase locus (UPRT; (39); kindly provided by David Sibley, Washington University) and a repair template consisting of 20 bp UPRT-targeting sequences flanking a PCR-amplified a dsRED expression cassette driven by the *T. gondii* GRA1 promoter with a GRA2 3’ UTR (amplified from the dsRED:LUC:BLEO plasmid described in (41, 42)) were used for transfection. Following centrifugation (800 × g, 10 min), 24 h *H. hammondi* zoites were resuspended in 450 μl of cytomix with ATP and glutathione containing 20 μg of the CRISPR/CAS9:*uprt*gRNA plasmid and 20 μg of PCR2.1 TOPO vector containing the dsRED repair template. The parasites were electroporated as above and each transfection contained at least 4 million parasites. Transfected parasites were then transferred to HFFs and grown for 2 days in cDMEM, then selected for 3 days by incubation in cDMEM containing 10 μM FUDR (Fig 5B). Parasites were again scraped, syringe lysed and filtered, and dsRED-expressing parasites were collected in PBS using flow cytometry. Sorted parasites were injected intraperitoneally into 2 BALB/c mice. After 3 weeks of infection, mice were euthanized, skinned, and the intestines were removed before feeding to specific pathogen-free cats. The oocysts were collected and purified as described above, and oocysts, sporozoites and replicating parasites were evaluated for dsRED fluorescence using microscopy, flow cytometry and FUDR resistance.

### FUDR resistance of transgenic parasites

To test for UPRT resistance in transgenic *H. hammondi*, sporozoites of wild type and dsRED-expressing parasites were incubated overnight in a T25 flask with confluent monolayer of HFFs. After 24 hours, parasites were isolated by needle passage and filtration as above. We infected coverslips containing confluent HFFs with 50,000 parasites, and parasites were exposed to media alone or media containing or 20 μM FUDR. The parasites were allowed to grow for 4 days and then fixed using 4%PFA for 20 minutes. After blocking overnight in PBS/BSA/triton, coverslips were stained with rabbit anti-*Toxoplasma* antibody at a 1:500 dilution for an hour, and stained with Alexa-fluor 488-labeled goat anti Rabbit antibody. For each treatment/strain combination, we counted the number of parasites in at least 100 vacuoles, and for HhdsRED parasites we also quantified growth in both red and wild type vacuoles (since the HhdsRED parasites were from a mixed population).

### RNA sequencing and data processing

Sporozoites were isolated from HhCatEth1 and TgVEG sporulated oocysts as described above and used to directly infect confluent HFF monolayers in 96 well tissue culture plates with MOIs of 3, 1 or 0.5 purified sporozoites for each species. On day 4 post-infection, wells were observed and chosen for RNA analysis based on similar numbers of parasite-containing vacuoles lack of significant host cell lysis in both species. For the Day 4 samples, the MOI=3 infected wells were chosen for *H. hammondi,* while the MOI=0.5-infected wells were chosen for *T. gondii*. Wells were washed 3X with ~200 μL cDMEM, and RNA was harvested using Trizol (Invitrogen). Remaining *H. hammondi* wells were washed with 3X with cDMEM on Day 4 and then again on Day 9 prior to harvest on Day 15 post-infection. For *T. gondii*, remaining MOI=3 and 1-infected wells were scraped and syringe lysed on Day 4 and Day 9 and used to infect new monolayers in 96 well plates at an MOI of either 0.5 (Day 4) or 0.3 (Day 9). On Day 15, wells for both *H. hammondi* (not subcultured) and *T. gondii* (subcultured 2x) were washed 1X with cDMEM and then harvested for RNAseq analysis. Two samples were harvested for each species on Day 4, and 3 samples were harvested from each species on Day 15. Total RNA samples were processed for Illumina next generation sequencing using the Mobioo strand-specific RNAseq library construction kit. Samples were analysed on an Illumina Nextseq and demultiplexed using NextSeq System Suite software. Reads were aligned to their respective species genome assembly (*H. hammondi* v10 or *T. gondii* strain ME49 v10; <u>toxodb.org</u>; (12)) using the Subread package for Linux (v. 1.4.6; (43)) with subread-align using default parameters except for –u to keep only uniquely mapping reads. featureCounts from the subread package (43) was used to quantify the number of mapping reads per transcript, using default settings except for –s 2 (for stranded read mapping), –t CDS, –g Parent (for specific compatibility with the *T. gondii and H. hammondi* gff files), –Q 10 (minimum mapping quality required). The –t CDS option was chosen for both *T. gondii* and *H. hammondi* because to date no 5’ or 3’ UTR sequences have been predicted for *H. hammondi* (in contrast to *T. gondii*). Fastq files have been deposited in the NCBI short read archive (Accession Numbers Pending).

### Identification of differentially expressed genes using DESeq2 and gene set enrichment analysis

Raw count data per transcript (generated by featureCounts above) were loaded into R statistical software and analyzed using the DESeq2 package (44). Comparisons of D4 and D15 read count data were used to identify transcripts of different abundance at each time point, and differences were deemed significant at *P*_adj_<0.05. Data were log_2_ transformed and normalized using the rlog function in DESeq2 for use in downstream analyses. Log_2_ transformed, normalized data from DESeq2 (hereafter referred to as Log_2_ (FPM)) were analyzed for enrichment using Gene Set Enrichment Analysis (GSEA; (24)). Since read count overall from the *T. gondii* libraries were much greater than those from the *H. hammondi* libraries, we took the 7372 genes matched based on the previously published gene-by-gene annotation (12) and selected only those that had at least 1 day 4 sample and 1 day 15 sample with >5 reads. In total, 4146 genes passed these benchmarks and were used in subsequent analyses. We used this approach to identify gene sets that were significantly different between days 4 and day 15 in culture in both species and those that were different between species at both day 4 and day 15. We compared these data using previously curated gene sets (434 total; as published in (25)) as well as 7 additional gene sets that we curated ourselves. These are listed in Table 2. All enrichment profiles were deemed significant if the FDR q-value was ≤ 0.05.

### cDNA synthesis and qPCR

For qPCR validation of RNAseq data, after 24 hours of growth, monolayers containing excysted sporozoites were scraped, syringe lysed, and pelleted. The pellet was resuspended in cDMEM, filtered through a 5 μM syringe-driven filter (Fisher Scientific, SLSV025LS) and passed at MOIs of 1 (*T. gondii*) and 7 (*H. hammondi*) into 96-well plates containing HFF monolayers. Mock-infected controls consisted of filtering parasites through a .22 μM syringe-driven filter (Fisher Scientific, SLGL0250S). Three replicates were made for each sample per isolation time point. RNA was collected at day 4 and day 15 post-infection, (equivalent to day 5 and day 16 post-excystation) from parasites and mock infections grown in 96-well plates. RNA was collected using the RNeasy Kit according to the manufacturer (Qiagen, 74104) using QiaShredder spin columns (Qiagen, 79654) to homogenize samples, and RNase-free DNase to degrade contaminating DNA (Qiagen, 79254). Isolated RNA was ethanol precipitated, resuspended in 10 ul RNase-free water, and these preparations were used to create cDNA using Superscript III Reverse Transcriptase Kit using Oligo(dT) primers according to the manufacturer (ThermoFisher Scientific, 18080051). All RNA samples were kept at -80°C, and all cDNA reactions were kept at -20°C. Prior to qPCR use, cDNA was diluted 1:10 with H_2_0.

qPCR assays were performed on a QuantStudio 3 Real-Time PCR System in a 10 μl reaction volume containing 5 μl 2x SYBR Green Master Mix (VWR International, 95030-216), 3 μl of cDNA template, 1 μl H_2_0, and 1 μl 5 μM primer. Controls included a reverse transcription negative control and a water-template control. The thermal cycling protocol was 95°C, 10 min; 40 cycles of (95°C, 15s; 60°C, 1 min); 4°C hold. The melt curve protocol was 95°C, 15 s; 60°C 1 min, 95°C 15 s. The control gene was dense granule 1 (GRA1), and samples were tested in duplicate or triplicate. Melt curves were performed on each plate (with the exception of 2 plates). Data were analyzed using the 2^-ΔΔCt^ method (45), and statistical analyses were conducted on the ΔCt values (as in (46)).

### Animal statement

All mouse experiments were performed with 4- to 8-wk-old BALB/C and C57BL/6J mice. Animal procedures met the standards of the American Veterinary Association and were approved locally under Institutional Animal Care and Use Committee protocol no. 12010130.

## Acknowledgements

The authors would like to thank Clara Stuligross for a critical reading of the manuscript. At the University of Pittsburgh, Cori Richards-Zawacki, Stephanie Ander, and Carolyn Coyne shared expertise, reagents, and equipment required for qPCR. This work was supported by R01AI116855 and R01AI114655 to J.P.B.

The authors declare no conflicts of interest.

## Supporting information

**S1 Fig.**
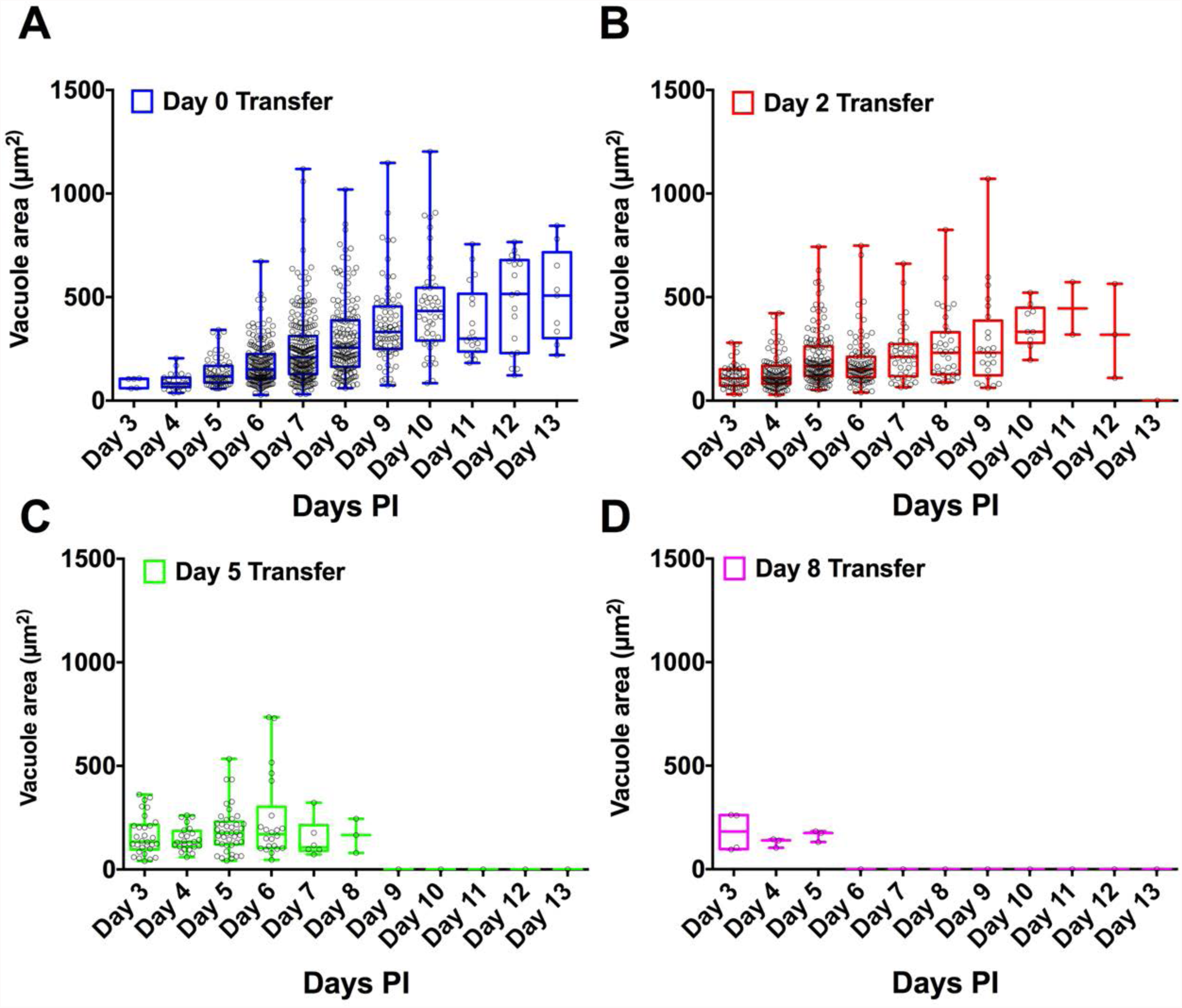
*H. hammondi* vacuole sizes following subculture. A-D) Visible vacuoles sizes were measured using ImageJ (NIH) for each day of attempted subculture. For each day when successful subculture was detected (Fig 5 C-H), vacuoles appear to grown in size, until they reach a plateau prior to the disappearance of vacuoles.

**S2 Fig.**
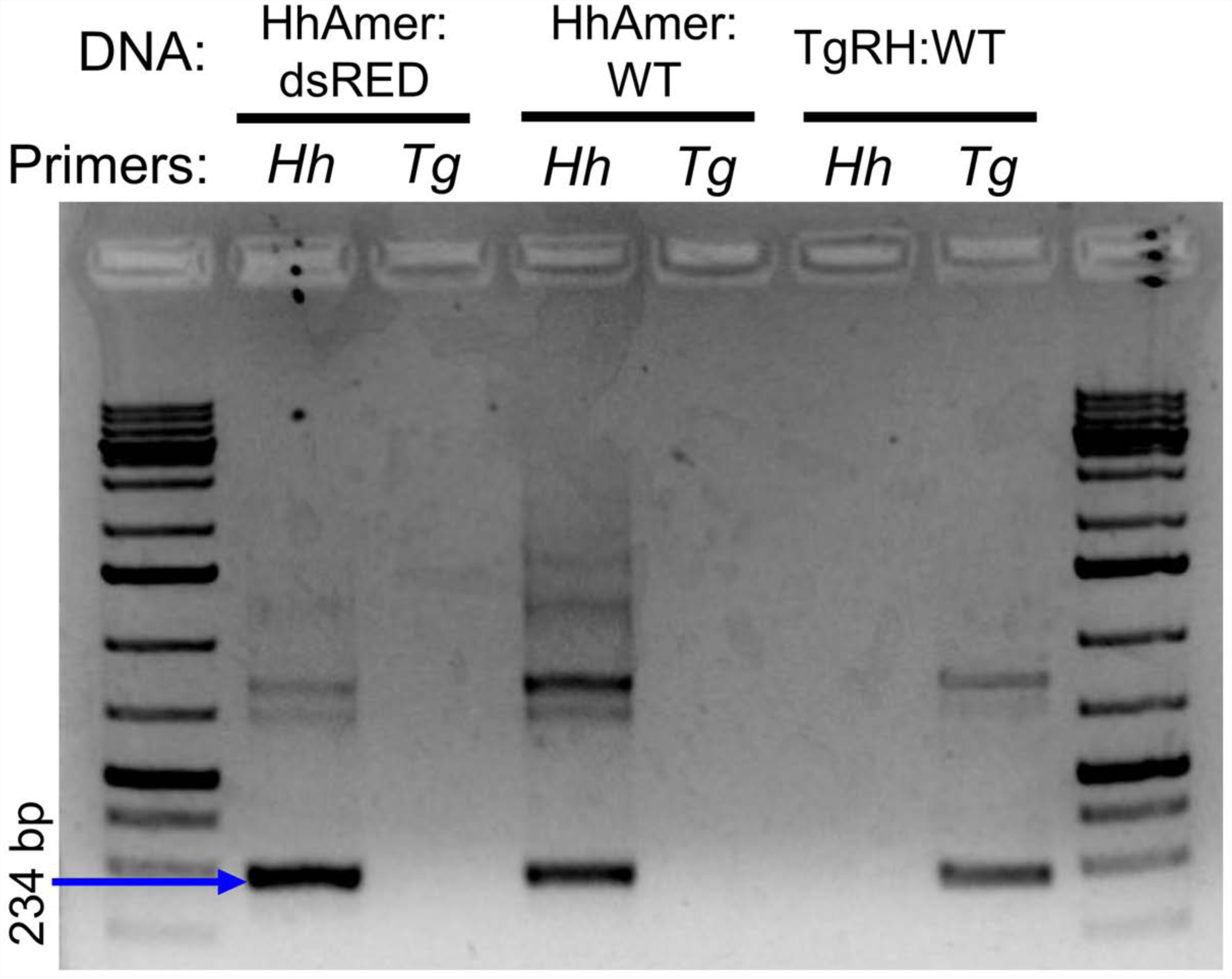
Validation of transgenic *H. hammondi.* PCR validation of *H. hammondi* dsRED-expressing parasites using *H. hammondi* and *T. gondii*-specific primers (see Table S2 for primer sequences).

**S3 Fig.**
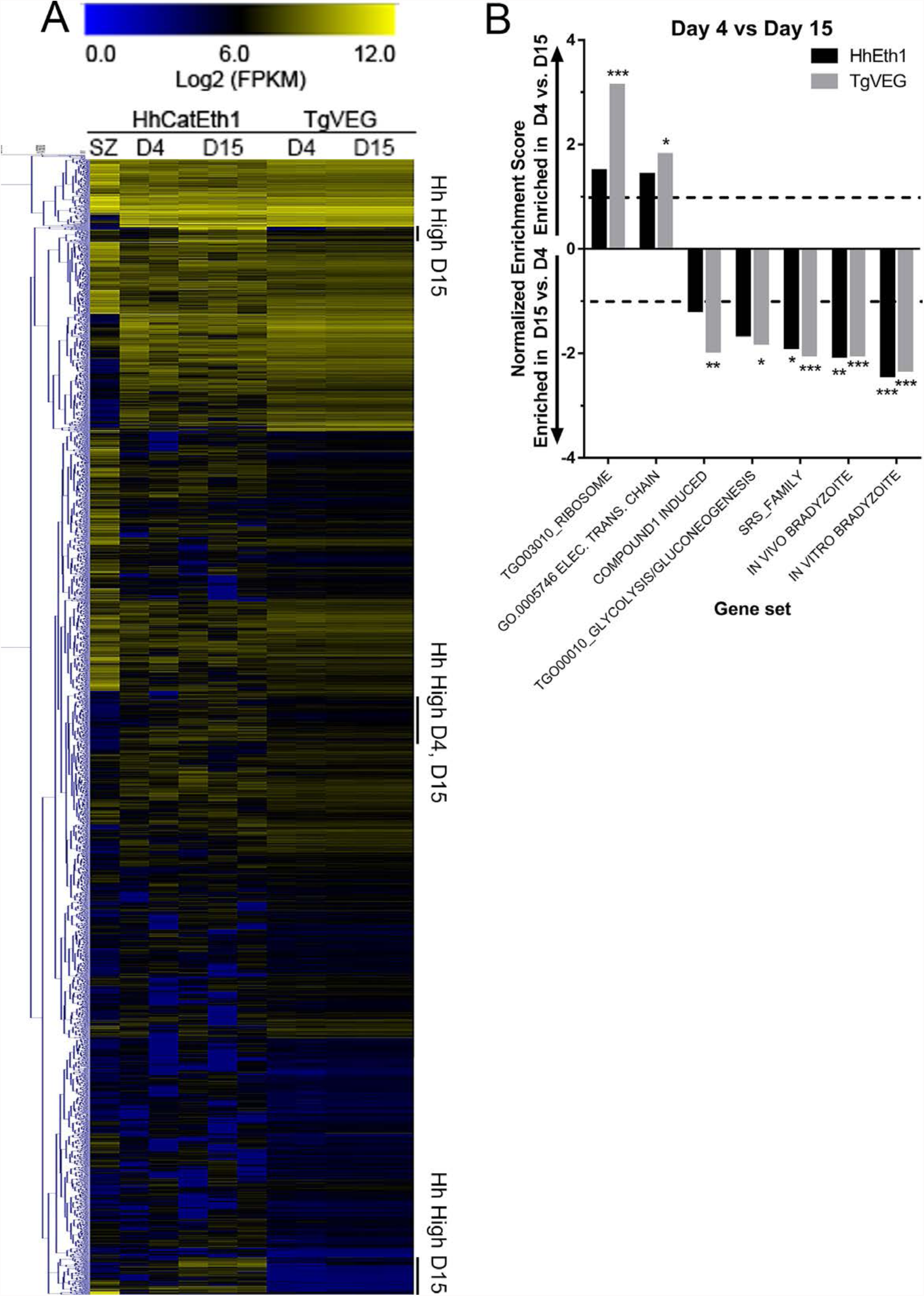
Transcriptional profiling and Gene Set Enrichment Analysis of *T. gondii* and *H. hammondi*. A) Hierarchical cluster (Euclidean distance, complete) of all 4276 genes with detectable expression in *T. gondii* and *H. hammondi*. Raw Log_2_ transformed FPM values (taken from DESeq2 following normalization and transformation using “rlog”) are shown. B) Gene set enrichment analysis results identifying gene sets that were significantly difference between D4 and D15 for each species. Gene sets are described in Table 2. *: P<0.05; **:P<0.01; ***:P<0.001.

**S4 Fig.**
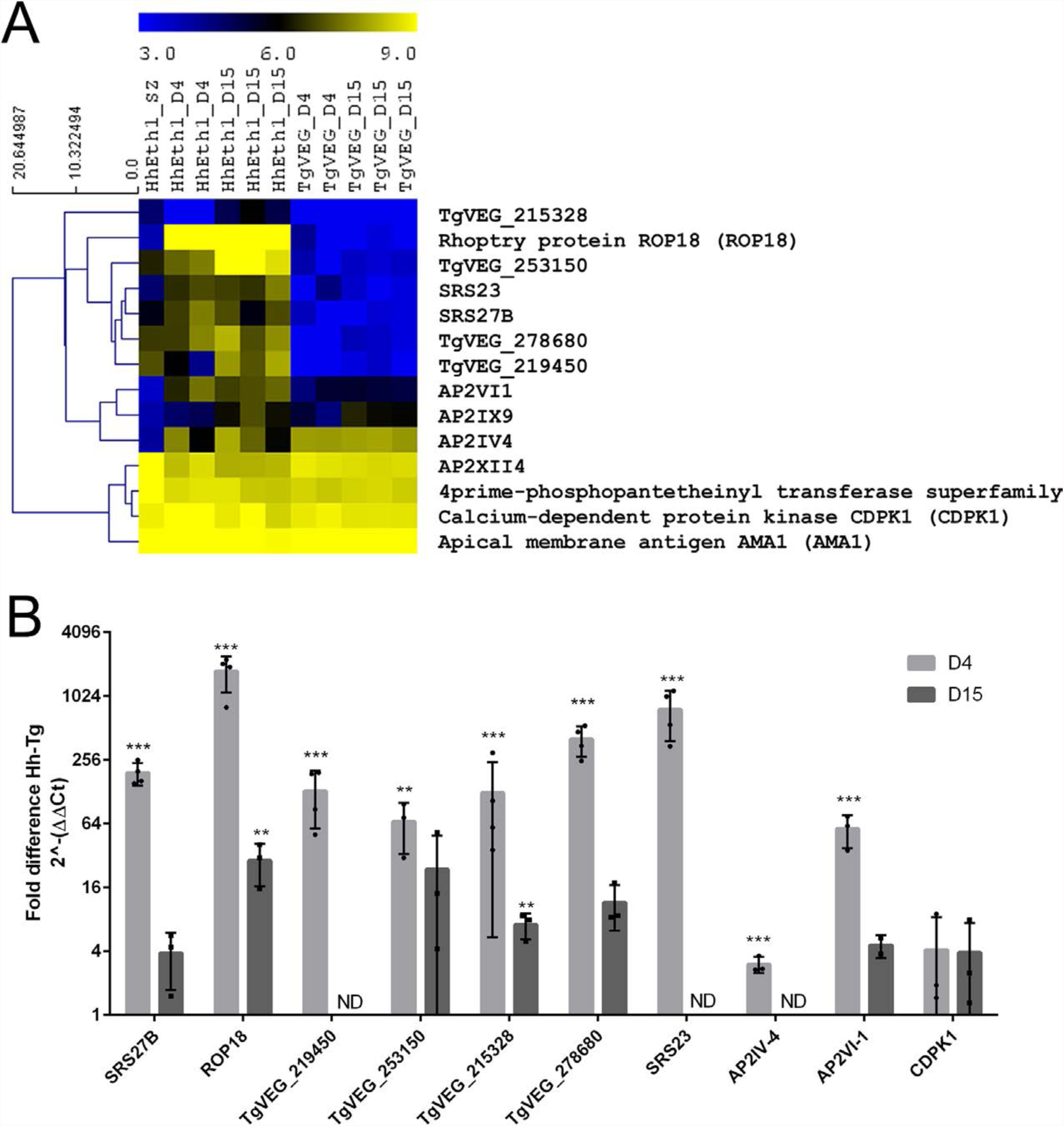
qPCR validation for 9 transcripts that were found to be of higher abundance in *H. hammondi* compared to *T. gondii*. A) RNAseq expression profile of HhCatEth1 and TgVEG at 4 and 15 DPI. B) qPCR validation of 9 transcripts demonstrate significantly higher transcript level in HhCatEth1 compared to TgVEG at day 4 (9 genes) and/or day 15 (2 genes) pi. Fold difference of HhCatEth1 genes relative to TgVEG is shown; bars represent mean and SD from three biological replicates. Significance determined from ΔCt values using multiple t-tests and the Holm-Sidak method, with alpha=5.0%. Calcium dependent protein kinase 1 (CDPK1) served as a control.

**Supplementary Table 2:**
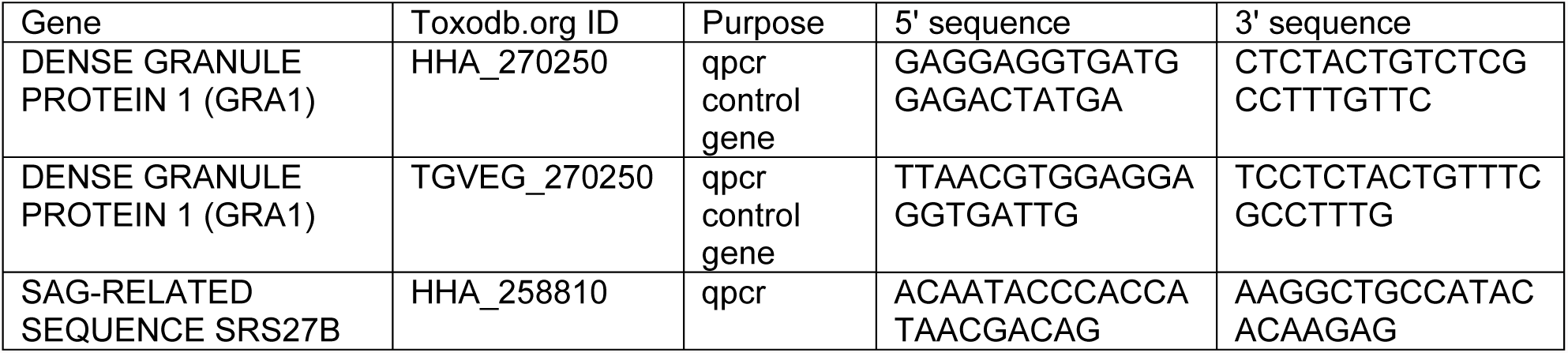
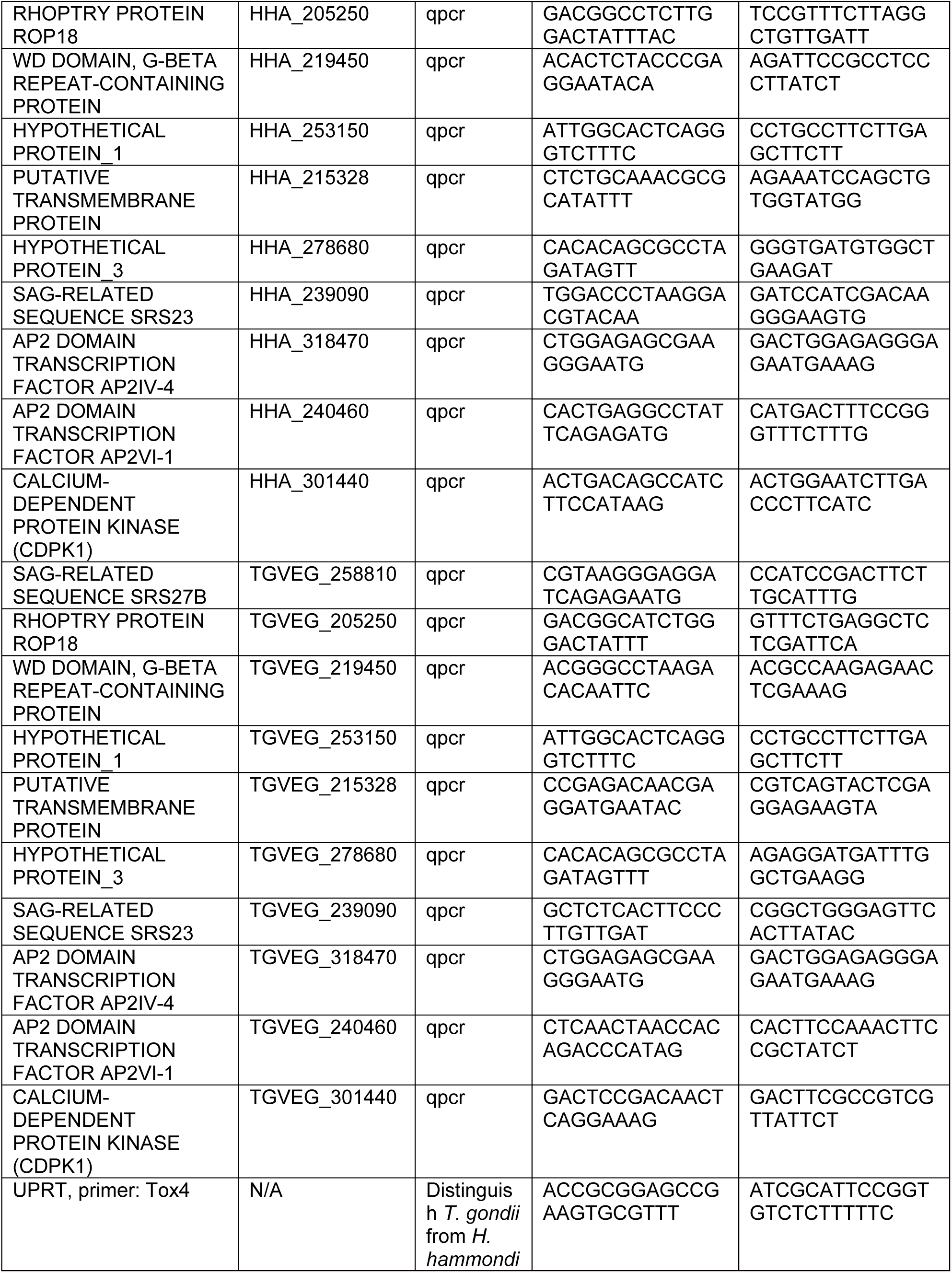
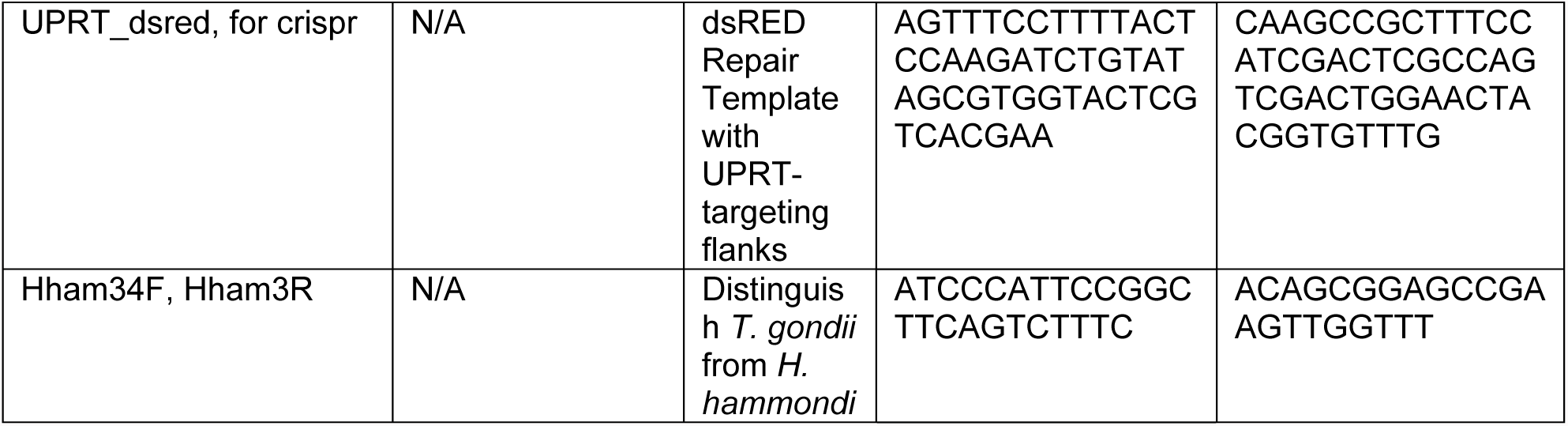
Primer sequences.

